# Gene Expression Reactomes Across Species Do Not Correlate with Gene Structural Similarity

**DOI:** 10.1101/2024.02.23.581748

**Authors:** Wei Chen, Yi Chai, Nancy Y. Cui, Ashwin Gopinath, David Yu Zhang

## Abstract

Mammalian animal model species for preclinical drug testing share a large number of genes with similar names and/or protein primary structures, but gene expression patterns in different tissues and in response to different drugs have not been previously systematically characterized. Here, we experimentally measure the gene expression responses (reactomes) of each gene in the mouse and rat genomes in 10 different organs to 3 different drugs. Surprisingly, we observe that gene reactomes across species generally do not correlate with structural similarity. Thus, we propose an alternative functional cross-species gene mapping approach, based on organ-specific gene expression response to drug dosing.

## Introduction

Drug testing in animal models is a critical and mandatory component of preclinical studies for drug development that lead to investigational new drug (IND) applications to regulatory agencies such as the Food and Drug Administration (FDA) or the European Medicines Agency (EMA). These animal studies are primarily focused on toxicology and safety [1], aiming to capture the systems biology effects of novel compounds on various organ systems. However, despite rigorous good laboratory practice (GLP) preclinical toxicology studies on animal models including non-human primates (NHPs), roughly 40% of new drug candidates will fail Phase I human clinical trials due to observations of severe adverse events in humans [2].

To study why safety/toxicology properties of drug compounds do not always effectively translate between even genetically similar mammalian species (NHPs and humans with >98% genetic homology [3]), we designed and performed systematic experiments to observe the impact of 3 small molecule drugs on the two most commonly used rodent animal model species, mouse (*mus musculus*) and rat (*rattus norvegicus*). Based on our previous high-frequency longitudinal transcriptomics study [4], we identified that (1) a significant number of genes’ expression levels are impacted by bleeding and drug injection, and (2) the largest number of genes’ expression levels are affected roughly 2 days after a single bolus dose of a small molecule drug. Consequently, we designed our experiments to include daily blood draws for 3 days to acclimate the animals to bleeding, dose the animals with drugs just after the Day 3 blood draw, and then sacrifice the animals after the Day 5 blood draw (Fig. 1a). Tissue samples from ten different organ systems were harvested following necropsy, and the RNAseq was performed on each of the samples. For each of 3 drugs and 1 saline control treatment, 3 male rats, 3 female rats, 3 male mice, and 3 female mice were used, for a total of N=48 animals and S=720 RNAseq samples.

**Figure 1.**
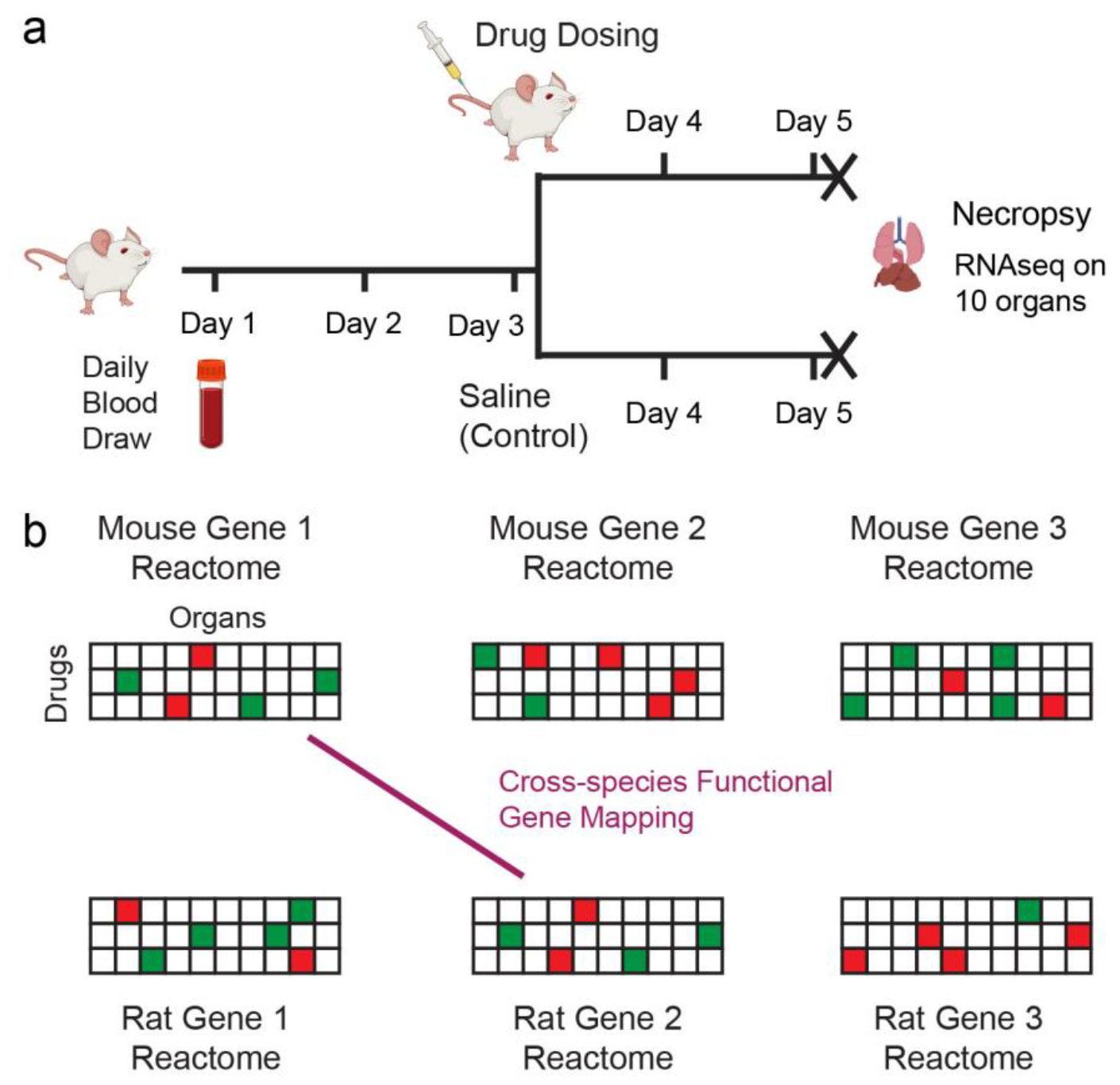
Overview of experimental design and cross-species gene mapping goal. **(a)** Blood was drawn daily from mouse (*mus musculus*) and rat (*rattus norvegicus*) models for 3 days, and then the animals were dosed with one of three drug molecules just after the Day 3 blood draw. Daily blood draws continued for 2 more days, before the animals were sacrificed on just after the Day 5 blood draw, and then necropsy was performed on 10 common organ systems. RNA extracted from the 10 different tissue samples, as well as the 5 daily blood samples, for a total of 15 RNAseq samples per animal. For each of 3 drugs and 1 saline control treatment, 3 male rats, 3 female rats, 3 male mice, and 3 female mice were used, for a total of N=48 animals and S=720 RNAseq samples. **(b)** For each gene in each species, we can construct a 30-dimensional scalar vector (3 drugs * 10 organs) corresponding to the gene’s expression change following drug dosing in each organ. We define this vector as the Gene Expression Reactome, or reactome for short. Genes across the two species can be functionally mapped to each other based on similarity of their reactomes.

One goal of this study is to characterize the Gene Expression Reactome (henceforth reactome) for each gene: the changes in its expression in each organ in response to each drug. In our current experiments, each gene’s reactome would comprise 30 values, with 1 scalar value corresponding to the mean gene expression change for each of 3 drugs * 10 organs. By comparing the reactomes of different genes across species, our goal is to be able to functionally map genes across species based on drug response (Fig. 1b). In the rat-mouse species pair, for example, there are 18,407 genes with the same name and high protein structural similarity, but more than 10,000 genes in each species with no clear corresponding gene in the other species. As the number of drugs and/or organs tested increases, the reactome becomes higher dimensional, allowing more precise gene mapping.

## Results

### Gene Expression Reactome Profiling and Results

Fig. 2a shows the normalized log2 expression of the rat *Bex1* gene using saline control injection in the 10 different organs as a radar plot, with individual traces shown for each of the 6 animals. The reproductive organ denotes testes for male animals and ovaries for female animals, and PBMC denotes the peripheral mononuclear blood cells separated from the blood buffy coat. See Methods and Supp. Section S1 for NGS data alignment and gene expression normalization details. For rat *Bex1*, some organs showed highly consistent expression levels across the 6 animals (e.g. brain), and other organs showed significant individual variability (e.g. kidney and heart). We believe that the variability of gene expression in control samples arises from a combination of the individual’s biological state and technical variability during the RNA extraction and next-generation sequencing (NGS) library preparation process, with the latter being a bigger contributor for low expression and the former being a bigger contributor at high expression. See Supp. Figs. S3-S6 for additional examples of control animal tissue gene expression for rats and mice.

**Figure 2.**
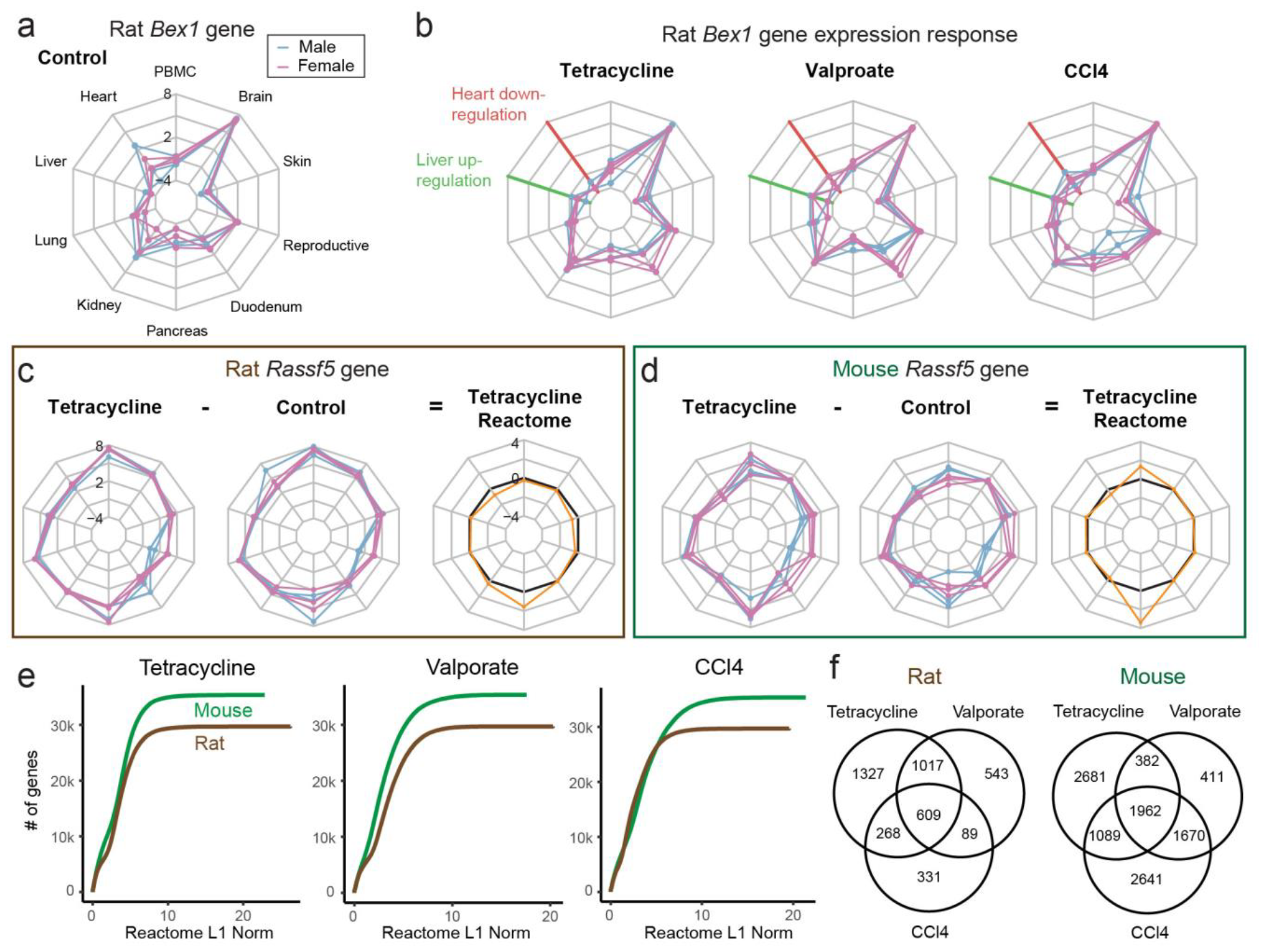
Example organ gene expression profiles and reactomes. **(a)** Gene expression for the rat *Bex1* gene in all 10 organs. All 3 male and 3 female animals’ expression levels are plotted individually to show variability. The plotted expression levels are in normalized log2 units; see Methods for bioinformatic pipeline details. **(b)** Expression of the rat *Bex1* gene after dosing with tetracycline, valproate, and carbon tetrachloride (CCl4). Up-regulation in the liver and down-regulation in the heart is observed for *Bex1* in response to dosing by all 3 drugs. This is not a general observation; the majority of genes showed reactomes with different responses to the 3 different drugs. **(c)** Illustration of the reactome computation for the rat *Rassf5* gene for tetracycline. **(d)** Illustration of the reactome computation for the mouse *Rassf5* gene for tetracycline. The mouse *Rassf5* gene has similar a tetracycline reactome as the rat *Rassf5* gene, though the mouse shows up-regulation in PBMC that is absent in rat. **(e)** Cumulative distribution function (CDF) plots of reactome L1 norm values for genes in the mouse and rat transcriptomes. Roughly 50% of genes exhibited reactome L1 norms below about 3 for all 3 drugs. **(f)** Venn diagram showing the number of genes significantly affected (greater than 2-fold change in expression change) in any organ by each of the 3 dosed drugs. The three drugs exhibited relatively distinct impact on the genes impacted, as seen by the significant populations of genes in each of the Venn diagram sectors.

Fig. 2b shows the rat *Bex1* gene expression in different organs after dosing with 200 mg/kg tetracycline, 500 mg/kg valproate, or 2000 mg/kg carbon tetrachloride (CCl4). The rat *Bex1* gene was selected for illustration here because it exhibits similar tissue gene expression responses for all 3 drugs: up-regulation in liver and down-regulation in heart. Most genes in general do not exhibit similar expression responses to all three drugs. See Supp. Figs. S7-S10 for additional examples of gene expression responses.

Fig. 2cd illustrates our calculation of the rat and mouse *Rassf5* gene’s tetracycline reactome values. The arithmetic mean of the normalized log2 expression values of all 6 animals for the control group are subtracted from the corresponding arithmetic mean of the normalized log2 expression values for the tetracycline dosed group to produce a single scalar number for each organ. Note that for the reproductive organ, the expression levels of the 3 male testes samples and the expression levels of the 3 female ovaries samples are averaged together. We chose to treat these reproductive organs as a single tissue group rather than 2 separate tissue groups because we observed that the vast majority of genes exhibit similar expression for ovaries and testes (see Supp. Section S2 and Supp. Figs. S11-12). See Supp. Figs. S13-15 for examples of gene reactomes for different mouse and rat genes to different drugs.

In this work, we use the L1 norm of the reactome to quantitate the overall degree of expression perturbation for a gene. The L1 norm is simply the sum of the absolute values of all scalar value components of the reactome. For example, the L1 norm of the vector (-1, 2, 0) is 3. Fig. 2e shows the cumulative distribution function (CDF) of the reactome L1 norm for all rat and mouse genes for tetracycline, valproate, and CCl4. From these plots, we see that there is a wide distribution of reactome L1 norm values for the mouse and rat genes, and a small number of genes with large reactome L1 norm values contributing to the long tail of the distribution. See Supp. Fig. S16 for partial distribution function (PDF) of the reactome L1 norm values for each drug. We chose to use the L1 norm metric because this analog approach avoids outsized changes in reactome values due to slight measurement errors around cutoff thresholds that would occur when discretizing gene expression changes (Supp. Figs. S17-19).

To assess the degree of overlap between genes affected by the drugs, we constructed lists of genes with gene expression change over 2-fold (1 units of normalized log2 expression) in any of the 10 organs (union) for each drug, and show their overlap distribution in Fig. 2f. For both mouse and rat, the genes impacted by the 3 drugs appear to be relatively independent, as all sectors of the Venn diagram exhibited a significant population of genes. This is a desirable outcome, become more independence between genes in their reactomes to different drugs allow more information for performing gene mapping. In an undesirable case where gene expression reactomes are nearly identical across the 3 drugs, the second and third drugs would not provide significant additional information for performing cross-species gene mapping.

### Protein Primary Structure Similarity Does not Correlate to Reactome Similarity

A total of 18,407 genes out of the 35,085 mouse genes and 29,376 rat genes share the same name, based on high protein structural homology. Given the close phylogenetic distance between mouse and rat, we expected that these same-name gene pairs across mice and rats would have relatively higher reactome similarity (low reactome L1 norm distance). Furthermore, the same-name gene pairs exhibit a distribution of protein structures, reflected in both the primary structure (amino acid sequence, Fig. 3a) and the AlphaFold folded structures (Supp. Fig. S20). Within these same-name gene pairs, we likewise expected higher reactome similarity in the gene pairs with higher structural similarity.

**Figure 3.**
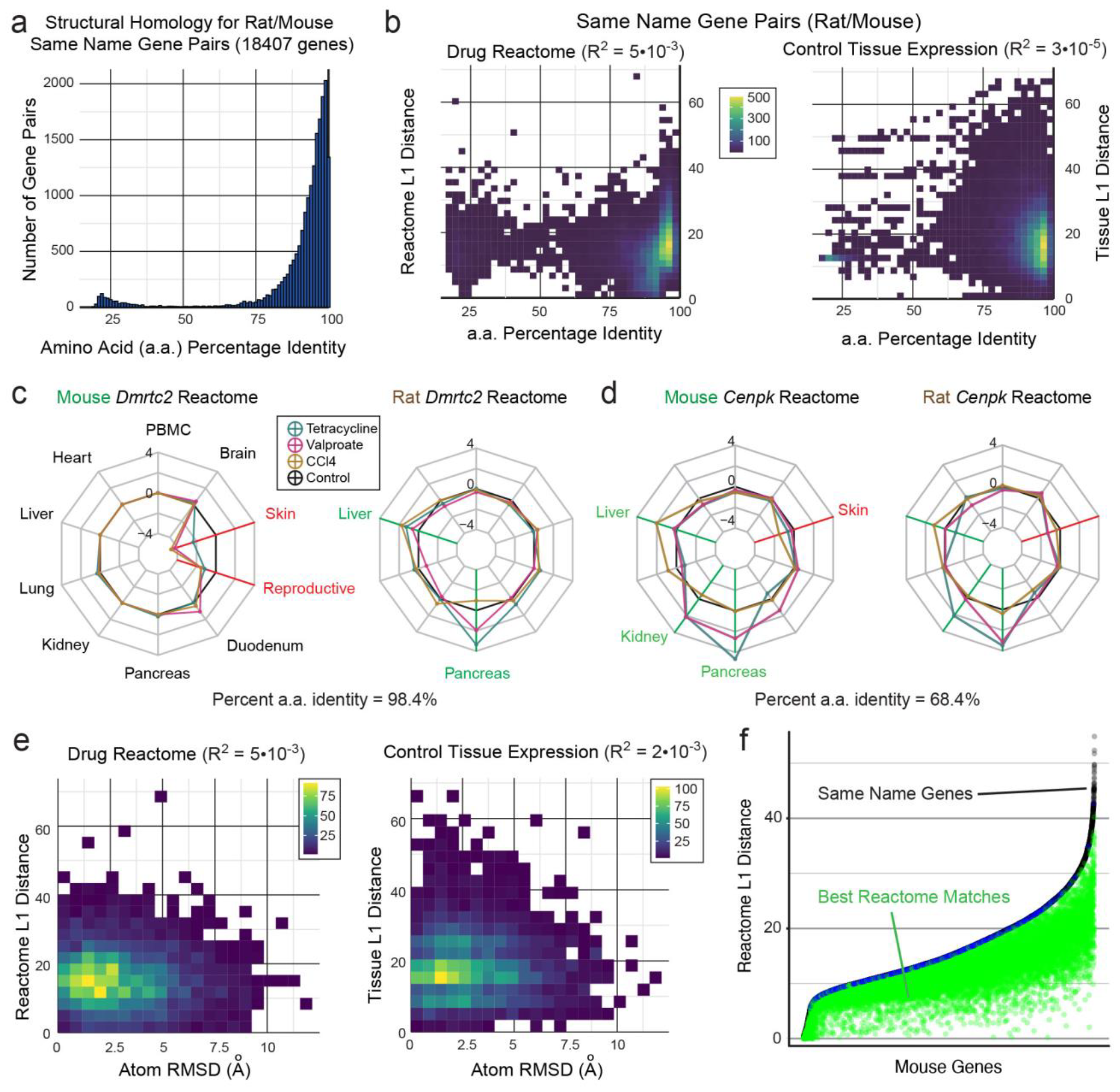
Gene pair reactome distance across rodent species does not correlate with gene structural similarity. **(a)** The rat and mouse transcriptomes share a total of 18,407 genes with the same names. These same-name gene pairs exhibit a range of protein primary structure similarities ranging from 20% to 100%, quantitated as the number of amino acid (a.a.) residues that are identical. **(b)** Control samples’ tissue gene expression L1 distance and drug Reactome L1 distance for same name gene-pairs are uncorrelated to a.a. identity between the genes. L1 distance denotes the sum of absolute values of log2 Expression difference between the two genes. This is an unexpected result; if protein structural similarity were correlated with drug response across species, then we would expect to observe a significant negative correlation (higher a.a. identity accompanies lower L1 distance). **(c)** *Dmrtc2* is an example of a same name gene pair with high a.a. identity and low Reactome similarity (high Reactome L1 distance). **(d)** *Cenpk* is an example of a same name gene pair with a low a.a. identity and high Reactome similarity. **(e)** Same-name gene pair structural difference, quantitated as protein atom root mean square distance (RMSD), also does not appear to correlate to reactome L1 distance or tissue expression L1 distance. **(f)** Analysis of reactome L1 distance for same name genes vs. best reactome matches. In only 400 same-pair genes (2.2%) were also the least reactome L1 distance match gene (blue).

Surprisingly, our data suggests <u>no significant correlation between structural similarity and reactome distance for same-name gene pairs (Fig. 3b)</u>. In particular, we note that <u>none</u> of the same-name gene pairs with over 98% a.a. identity exhibited reactome L1 distance of below 2 (the lowest bin). Furthermore, the gene expression of same-name gene pairs across organs in the control animals also showed essentially no correlation with the structural similarity. These findings indicate that organ-specific gene expression, both at homeostasis and in response to drug dosing, do not translate across even closely related species. By extension, to the extent that toxicology of new drug molecules are correlated across species, they may manifest in very different ways affecting different sets of genes. The details of gene regulatory networks in systems biology appear to be species-specific.

Given the unexpected nature of our findings, we carefully analyzed the detailed reactomes of a number of same-name genes to spot-check our conclusions. Fig. 3c shows the reactomes of the rat and mouse *Dmtrc2* gene (98.4% a.a. identity); the responses are highly distinct with mouse *Dmtrc2* down-regulated in response to all 3 drugs in skin and reproductive organs, and rat *Dmtrc2* up-regulated in live and pancreas. Fig. 3d shows the highly similar reactomes of the rat and mouse *Cenpk* gene (68.4% a.a. identity). See Supp. Figs. S21-24 for additional examples of same-name gene pair reactomes.

To verify that the lack of correlation between protein amino acid sequence and drug response reactome also reflects a lack of correlation between protein 3-dimensional structure and drug response reactome, we next analyzed the Alphafold-predicted structures of same-name gene pairs between mouse and rat. As expected, the root-mean-square distance (RMSD) for atoms in the same-name gene pairs, a commonly accepted measure of protein structural similarity with lower RMSD indicating higher structural similarity, also has essentially no statistical correlation with the reactome L1 distance (Fig. 3e).

Finally, we analyzed the mouse-rat gene pairs with the lowest reactome L1 distances (highest reactome similarity), to check if there is correlation between reactome L1 distance and protein a.a. identity in this group (Supp. Fig. S25). In certain unusual statistical distributions, one may observe correlation in some subsets but not others or the superset (see Simpson’s Paradox). Consistent with our earlier findings, we observe no correlation for higher protein a.a. identity based on the best-match gene pairs based on reactome L1 distance. In only 400 out of the 18,407 genes (2.1%) were the same-name gene also the best reactome match (Fig. 3f).

### Cross-Species Functional Gene Mapping based on Reactome

Given our observation that some cross-species gene pairs exhibit significantly better reactome similarity than both same-name gene pairs, we next explore the functional mapping of genes across species based on the reactome. We quantitate the degree of gene matching using Shannon Entropy (Fig. 4a), a concept commonly used in information theory to represent the degree of uncertainty of a variable.

**Figure 4.**
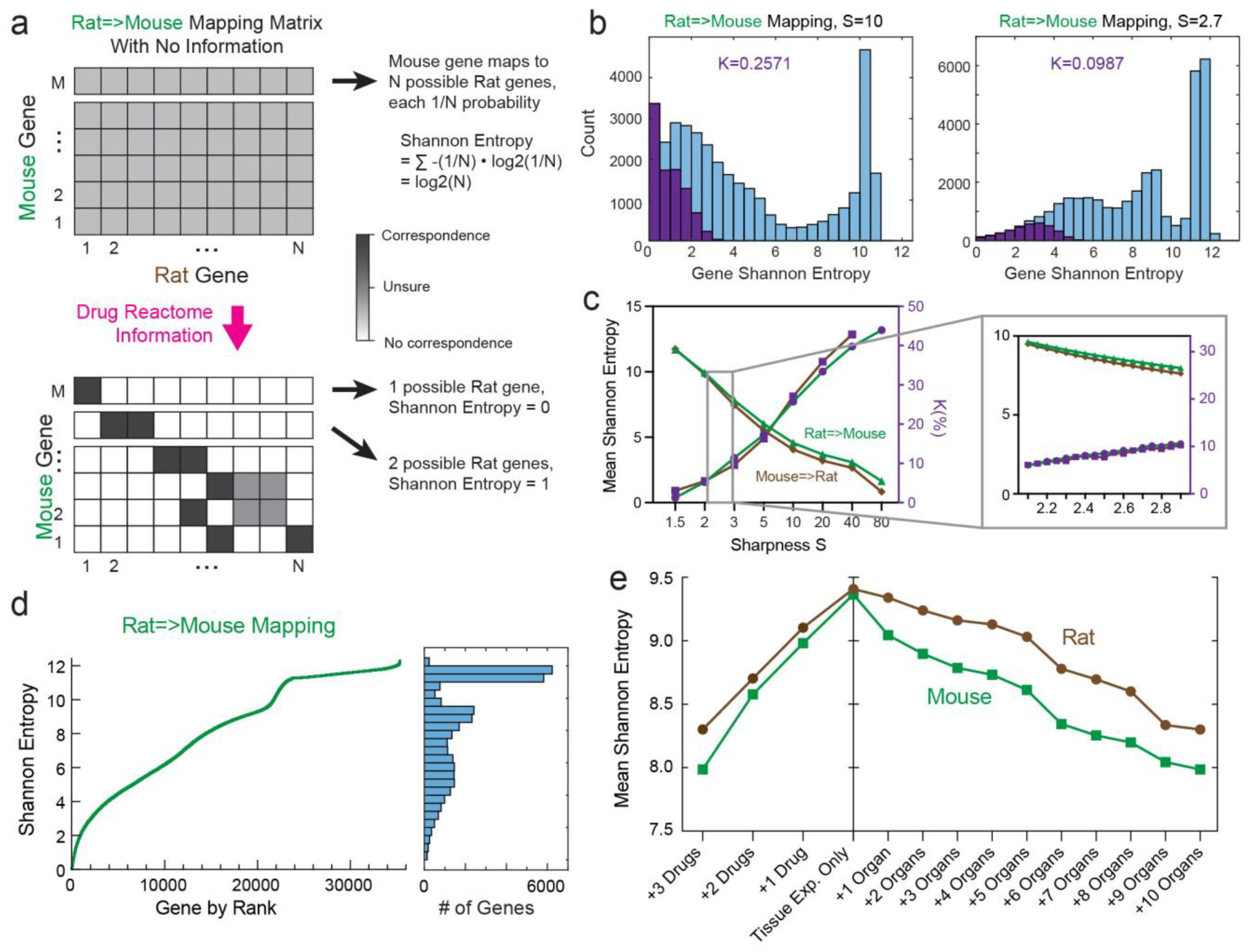
Shannon Entropy of cross-species functional gene mapping based on reactomes. **(a)** Explanation of Shannon Entropy in the context of cross-species gene mapping. **(b)** Impact of sharpness S on the Shannon Entropy distribution and the maximum false matching rate K. The purple histograms show the Shannon Entropy distribution for a “shuffled” dataset in which the 35,085 mouse genes’ reactome values for each organ/drug pair is randomly permuted, with frequencies scaled by K. At the relatively high value of S=10, K = 0.2571 (left) for rat=>mouse gene mapping. A similar Shannon Entropy distribution and value of K are observed for mouse=>rat gene mapping (Supp. Fig. S29) The right panel shows the Shannon Entropy distribution for S=2.7, the maximum value of S for which K < 0.10 for both rat=>mouse and mouse=>rat gene mapping. **(c)** Dependence of mean Shannon Entropy and K for different values of S. For the remainder of the figure, S=2.7 was used. **(d)** Distribution of Shannon Entropies for mouse genes, using both control tissue expression and drug reactomes, sorted in ascending order. The histogram of the Shannon Entropies to the right correspond to the right panel of subfigure (b). **(e)** Scaling properties of Shannon Entropy as additional drug reactome data or additional organ reactome data are included. The mean Shannon Entropy of gene mapping decreases by about 0.4 to 0.5 per drug, and by about 0.12 to 0.15 per organ.

In mapping the N possible rat genes to one mouse gene, initially we start with no information, so every rat gene is equally likely to be mapped and the Shannon Entropy on the mouse gene is computed to be log2(N). The Shannon Entropy for the two species is asymmetrical; each rat gene starts with Shannon Entropy of log2(M). As rat genes are excluded from matching to a particular mouse gene based on the reactome values, the Shannon Entropy of the mouse gene decreases to a minimum of 0 (perfect matching). In practice, we do not expect any gene pairs to reach 0 Shannon Entropy because of measurement errors on expression and because of differences in biology.

Briefly, we calculate the Shannon Entropy E_mouse_(i) of mouse gene i based on the following formulas:

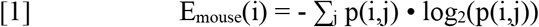

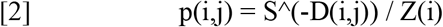

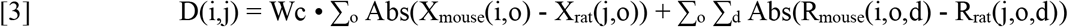

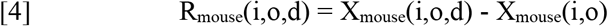

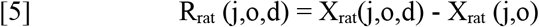

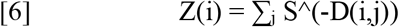

p(i,j) denotes probability of a match between mouse gene i and rat gene j

S denotes the sharpness of the probability dependence on reactome distance

D(i,j) denotes the reactome distance between mouse gene i and rat gene j

Wc denotes the control tissue expression weighting

X_mouse_(i,o) denotes the expression (normalized log2 units) of mouse gene i in organ o

X_rat_(j,o) denotes the expression of rat gene j in organ o

R_mouse_(i,o,d) denotes the reactome (change in expression) of mouse gene i in organ o for drug d

R_rat_(j,o,d) denotes the reactome (change in expression) of rat gene j in organ o for drug d

Z(i) denotes the partition function of the mouse gene i

Our Shannon Entropy calculations depend sensitively on two hyper-parameters: control expression weighting W_C_, and probability sharpness S. The value of Wc describes the relative weighting of similarity of expression levels in control animals vs. drug response reactomes. The value of sharpness S describes how strongly the method favors a marginally better match (in the form of lower reactome L1 distance).

In general, larger values of S lead to smaller values of Shannon Entropy E, but increases the rate of false matching between unrelated genes. In the extreme case of S = infinity, then the method would assign probability 1 to matching between a rat gene and a mouse gene with the lowest reactome L1 distance, even if there’s a second mouse gene that is nearly identical in its reactome (e.g. off by 0.01). In practice, two mouse genes with nearly identical reactomes (i.e. within RNA expression measurement noise) should both be assigned probability 0.5, assuming there are no other mouse genes that have remotely similar reactomes. In contrast at the other extreme case of S = 1, then the method essentially ignores all information provides by the reactome, and assigns equal probability to all mouse genes to map to a particular rat gene. Consequently an intermediate value of S is ideal to balance the sensitivity and specificity of cross-species gene matching, with higher values of S favoring higher sensitivity but yielding lower specificity.

To quantitate the degree of potential false cross-species gene mapping, we created a “shuffled” dataset in which the 35,085 mouse genes’ reactome values for each organ/drug pair is randomly permuted. For example, in the shuffled dataset, gene M1 may be randomly assigned the liver•tetracycline reactome value for M2, the liver•valproate reactome value for M2000, the lung•CCl4 reactome value for M12000, etc. In this shuffled dataset, any gene mapping between the rat and mouse transcriptomes would be purely coincidental, and all gene mapping would by construction by nonspecific.

For a particular sharpness value S, we can generate a Shannon Entropy E histogram distribution for the shuffled dataset. In general, this E distribution will be biased to lower values of E, compared to the real dataset. We define K as the maximum value of the scaling factor on the E histogram for the shuffled dataset that allows the scaled shuffled dataset E histogram to be circumscribed by the biological dataset E histogram. In Fig. 4b, the purple bars shows the shuffled dataset E histogram scaled by K and the blue bars show the biological dataset Shannon Entropy histogram for based on the real data.

One way of interpreting the value of K is the maximum false matching rate. In other words, (1-K) is the minimum specificity of the method for a given value of S. At S = infinity, the observed value of K approaches 1, but presumably a large fraction (e.g. at least 50%) of the mapped gene pairs with minimal least reactome distance are actually correct. We find that S = 2.7 is the maximum value of S that ensures K < 10%, corresponding to at least 90% specificity (Fig. 4c), and use S=2.7 for the remaining analysis. See Supp. Figs. S26-28 for additional analyses of the effect of sharpness S. Fig. 4d shows the Shannon Entropy distribution of mapping rat genes to mouse genes for S=2.7.

One powerful feature of our approach to cross-species gene mapping based on reactomes is that the approach is scalable. As we increase the amount of expression data collected in response to additional drugs and/or from additional organs, we increase the dimensionality of the reactome vectors. Fig. 4d shows the mean Shannon Entropies of mouse and rat genes, starting from just considering control tissue expression levels, and successively adding the reactomes for each drug. We observe a steady trend of declining Shannon Entropies with data from each additional drug, with a marginal decrease of 0.4 to 0.5 units of Shannon Entropy per additional drug. If this scaling law holds with additional drugs, we expect to minimize the Shannon Entropy of cross-species gene mapping to near 0 with 16 to 21 additional drugs (for 19 to 24 in total). Critical to this assumption of continue linearity in Shannon Entropy decrease is that the additional drugs must be relatively independent in their mechanisms of action. For the 3 drugs tested here, Fig. 2f shows that they are relatively independent in terms of the genes that they affect.

Fig. 4d also shows the scaling of the Shannon Entropy based on increasing the number of organs reactomes analyzed. Like with drugs, there is a consistent decrease in Shannon Entropy as additional tissue types are analyzed. The current organs included already cover the most common organ systems, but these organs/tissue types could be further subdivided. For example, brain tissue could be subdivided by lobe, and small intestines could include ileum and jejunum tissue in addition to duodenum. Given that one important application of gene mapping is to predict toxicology effects for new potential drug molecules, however, we believe that scaling via additional drugs would be preferable to additional tissue types.

### Whole Blood Reactome

In addition to the endpoint organ samples, we also performed RNAseq on whole blood collected on a daily basis from each animal. We note that whole blood is distinct from the PBMC sample types that we analyzed for reactomes, because roughly 98% of the whole blood RNA derives from red blood cells, and PBMCs comprise only a small fraction of the remaining 2% of cells in the buffy coat. The longitudinal samples collected and analyzed are whole blood because only 20uL of whole blood could be collected from mice on a daily basis without severely affect the animals’ health. Separating PBMCs from 20uL of whole blood is not currently feasible with any commercially available instruments or solutions.

Our previous study on longitudinal whole blood RNAseq analysis [4] indicated that the whole blood RNA expression contained significant number of temporally varying genes (TVGs) that generate expression response to small molecule drug dosing. Here, we compare the whole blood expression response to that of the endpoint organ responses (Fig. 5abcd). We find that whole blood RNA expression correlate highest with PBMC and pancreas. However, a significant fraction of genes (10%-30%) with expression responses in whole blood are not reflected in any other organ (Fig. 5ef). Considered in this way, whole blood (and by extension red blood cells) can be potentially considered as a different organ/tissue type, with its own pattern of drug responses.

**Figure 5.**
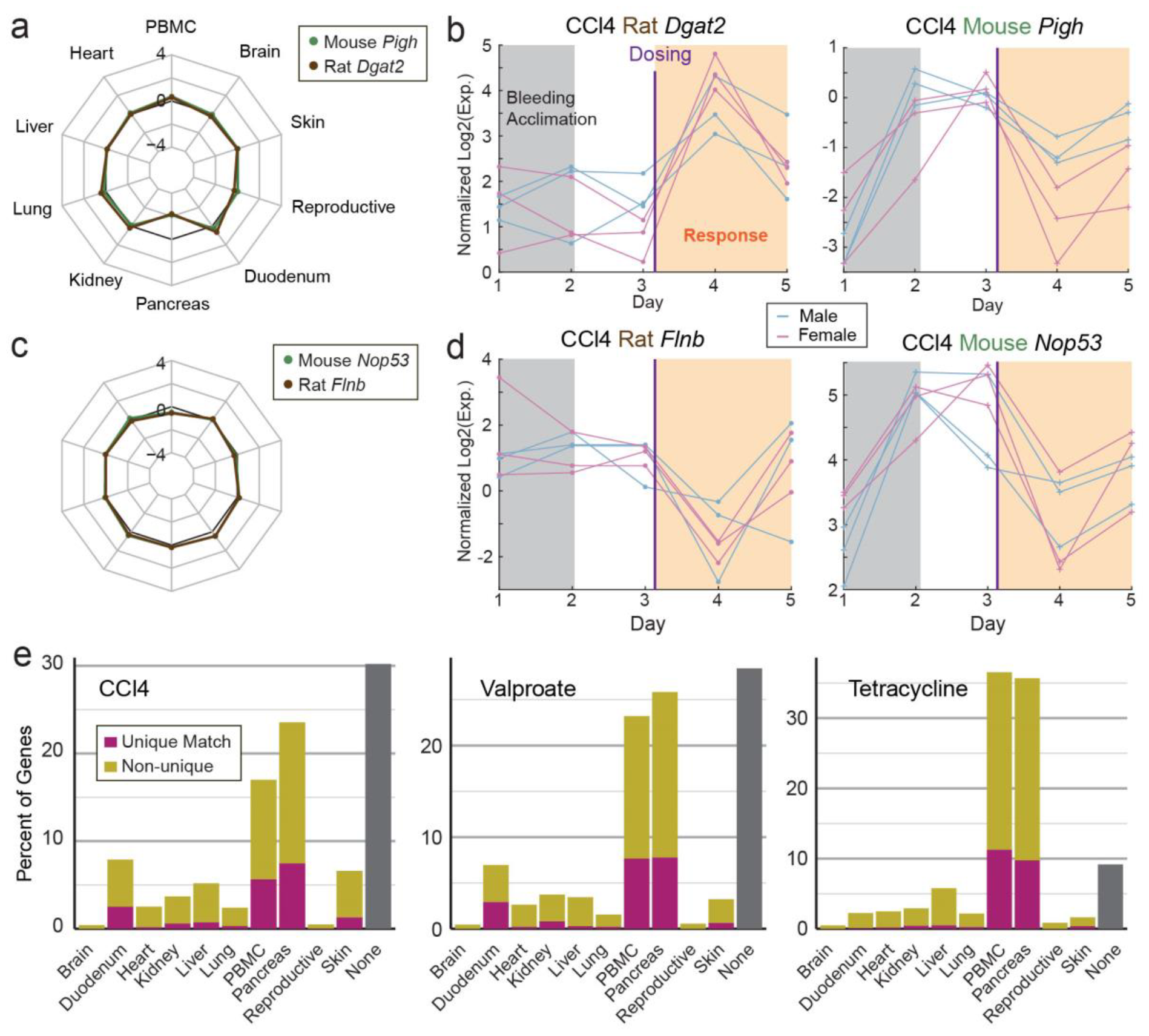
Whole blood longitudinal reactome vs. organ endpoint reactome. **(a)** The mouse gene *Pigh* and the rat gene *Dgat2* show highly similar endpoint organ reactomes to CCl4. **(b)** The temporal expression patterns in whole blood of mouse *Pigh* and rat *Dgat2* appear to be very different from each other, with rat *Dgat2* shows up-regulation response and mouse *Pigh* showing down-regulation response. Furthermore, mouse *Pigh* exhibits a bleeding acclimation up-regulation response on Day 2 that is absent in rat *Dgat2*. **(c)** The mouse *Nop53* and rat *Flnb* genes are observed to have no expression response to CCl4 for all organ endpoints. **(d)** In temporal whole blood samples, both genes are observed to have significant down-regulation response. The mouse *Nop53* further exhibits an up-regulation bleeding acclimation response. **(e)** Summary of gene expression reactome response in temporal blood samples vs. their responses in different organs at endpoint. Whole blood reactome does not appear to correlate strongly with any of the organs we characterized. Pancreas and PBMC showed the highest overlap in drug response genes with whole blood, but in both cases the overlap was less than 40% of genes for all 3 drugs.

## Discussion

In the course of systematic characterization of multi-organ gene expression responses to small molecular drug dosing across 2 rodent species, our primary finding is the counter-intuitive observation that rat and mouse gene pairs with high protein structural homology do not appear to exhibit high similarity gene expression reactomes. Conversely, the rat and mouse gene pairs that closest resemble each other from a drug response perspective also do not appear to associate with higher primary structure homology. One implication of this finding is that, while structural predictions and characterization of proteins may be useful for predicting drug efficacy within a species, protein structure prediction is generally less useful for systemic toxicity predictions, particularly when applied to cross-species translational medicine.

The biological basis for this finding will require more detailed studies to fully understand, but one hypothesis for explaining this observation may be that gene reaction networks and genetic regulatory elements are under smaller evolutionary pressure than primary amino acid sequence, and thus more prone to genetic drift on the timescale of evolution. By way of analogy, the Phillips-head screw has been in use in relatively unchanged form for more than 90 years after its initial patent, but the instrument and ways in which the Phillips-head screw is being used today is unimaginable to its inventors from the 1930’s. Similarly, biology and evolution appears to repurpose gene/protein components divergently in different organisms, resulting in different gene expression responses to drugs. By extension, this could partially explain why new drug toxicology predictions is so challenging and result in a 40% failure rate of Phase I human clinical trials.

Based on the above observation, we propose a new method to functionally map genes across species via the similarity of their reactome vectors. In addition for cross-species gene mapping, potentially for diagnostic biomarker discovery purposes, the reactome vector could also become a powerful tool for artificial intelligence (AI) methods. Specifically, the reactome vector, either in original analog format or in a digitized format following thresholding, could serve as a set of conditional tokens in the input of a generative AI based on transformers. This could allow the development of a pan-species foundation AI model that deeply understands systems biology and could perform inference on multiple species.

