## Supplementary Information for "Gene Expression Reactomes Across Species Do Not Correlate with Gene Structural Similarity"

### Methods

#### 1. Animal Study Design

**Table 1** Animal study design

| Group | Species & Strain | Age | Gender | Quantity | Compound Treatment | Dose Level | Dose Day | Sacrifice Day |
|---|---|---|---|---|---|---|---|---|
| 1 | SD Rat | 8 wks | M | 3 | Sterile NaCl | - | - | Day 5 |
| | | | F | 3 | | | | |
| | C57BL/6 Mouse | 8 wks | M | 3 | | | | |
| | | | F | 3 | | | | |
| 2 | SD Rat | 8 wks | M | 3 | Tetracycline | 200 mg/kg | Day 3 | |
| | | | F | 3 | | | | |
| | C57BL/6 Mouse | 8 wks | M | 3 | | | | |
| | | | F | 3 | | | | |
| 3 | SD Rat | 8 wks | M | 3 | Valproate | 250 mg/kg | | |
| | | | F | 3 | | | | |
| | C57BL/6 Mouse | 8 wks | M | 3 | | | | |
| | | | F | 3 | | | | |
| 4 | SD Rat | 8 wks | M | 3 | $CCl_4$ | 2 ml/kg | | |
| | | | F | 3 | | | | |
| | C57BL/6 Mouse | 8 wks | M | 3 | | | | |
| | | | F | 3 | | | | |
| 5 | SD Rat | 8 wks | M | 3 | Isoniazid | 300 mg/kg | | |
| | | | F | 3 | | | | |
| | C57BL/6 Mouse | 8 wks | M | 3 | | | | |
| | | | F | 3 | | | | |

Animal experiments in this study were approved by the Animal Care Committee at Biostate.AI (approval number: 101) and HD Biosciences Co. Ltd (approval number: 118-8). Sprague-Dawley rats were purchased from certified provider - Charles River Laboratories International, Inc (Shanghai, China). Rats were acclimatized prior to the experiment and were housed in enriched and ventilated housing cages throughout the experimental phase. Cage litter was changed at least once a week. The rats were housed under specific-pathogen-free (SPF) condition and subjected to a normal 12 hours light and night cycle, at 22 ± 2°C and 50 ± 10 % relative humidity. Chow and water were available *ad libitum*. The health conditions were examined and recorded daily by a veterinarian.

15 male SD rats and 15 female SD rats were housed for 5 days with daily 200 uL blood extraction. 15 male C57BL/6 Mouse and 15 female C57BL/6 Mouse were housed for 5 days with daily 50 uL mL blood extraction. Rats and mice in group 1 (control) were intraperitoneally injected with 0.9% sodium chloride on day 3. Rats and mice in the group 2 were intraperitoneally injected with tetracycline dissolved in sterile saline on day 3. Rats and mice in the group 3 were intraperitoneally injected with Valproic acid diluted in sterile saline on day 3. Rats and mice in the group 4 were administered a 1:1 mixture of corn or olive oil and carbon tetrachloride orally on day 3. Rats and mice in the group 5 were intraperitoneally injected with Isoniazid dissolved in sterile saline on day 3. All rats and mice were sacrificed on Day 5.

Detailed animal study design is shown in Table 1.

### 2. Bio samples collection and storage

200 μL of whole blood was collected daily from the same site of jugular vein of each rat between 10:00 am and 10:29 am. 50 μL of whole blood was collected daily from the same site of jugular vein of each mouse between 10:00 am and 10:29 am.

Whole blood samples were immediately processed for RNA extraction. On the final day of the study, rats were fasted overnight and then euthanized by cervical dislocation. Whole blood was immediately collected via heart puncture under RNase-free conditions. 1 mL of whole blood underwent the standard PBMC separation process, and the rest was preserved at -80°C for RNA extraction.

On the end day of the study, liver, kidneys (left and right), lung, heart, back skin, duodenum, brain, pancreas and reproductive (testis for male and ovary for female) were collected and underwent standard tissue RNA extraction process. The rest were preserved at -80°C.

### 3. RNA extraction from whole blood samples and liver samples

For RNA extraction from whole blood, 100 uL of whole blood was mixed with 700 uL of Trizol reagent and then underwent standard TRIzol and Chloroform RNA extraction method. Extracted RNA was further purified using Automatic Nucleotide Isolation Machine (Cat# NPA-32P, Bioer Technology, China) with MagaBio Plus Total RNA Purification Kit (Cat# BSC53M1B, Bioer Technology, China).

For RNA extraction from liver samples, around 30 mg of chopped liver tissue was mixed with 700 uL of TriZol Reagent and Lysing MatrixD. The mixture was ground for 2 min and then mixed with 140 uL of Chloroform. After 2 min of incubation at room temperature and 10 min of centrifugation at 12000 rpm and 4°C, supernatant was transferred to a tube and was further purified with MagaBio Plus Total RNA Purification Kit.

Purified RNA was quantified by using Nanodrop 2000 (Thermo Fisher, USA). The quality of RNA was measured by using RNA ScreenTape Assay (Agilent, USA) and 4200 TapeStation System (Agilent, USA).

### 4. mRNA capture and Release

VAHTS mRNA Capture Beads (Cat# N401-02, Vazyme Biotech, China) was used for the enrichment of mRNA from total RNA extracted from blood and tissue samples. During the mRNA capture and release process, mRNA was fragmented.

### 5. cDNA Synthesis and Library Preparation

Library preparation of mRNA including three steps: reverse transcription, adaptor ligation, index PCR amplification. Reverse transcription and adaptor ligation were performed using VAHTS Universal V6 RNA-seq Library Prep Kit for Illumina (Cat# N604-2, Vazyme Biotech, China) with standard protocol. Index PCR amplification was performed using VAHTS RNA Multiplex Oligos Set 1 for Illumina (Cat# N323-01, Vazyme Biotech, China) and VAHTS RNA Multiplex Oligos Set 2 for Illumina (Cat# N324-01, Vazyme Biotech, China) with standard protocol. One round of 1.6 × beads (Cat# CNGS-0500, CleanNA, Netherlands) purification was then performed after adaptor ligation. One round of 1.6 × beads purification was performed after index PCR amplification. Library DNA were quantified by Qubit 4.0 using Equalbit 1 × dsDNA HS Assay Kit (Cat# EQ121-01, Vazyme Biotech, China). Library was analyzed by using D1000 ScreenTape Assay (Agilent, USA) and 4200 TapeStation System.

### 6. Sequencing

2 ×150 paired-end sequencing was performed on Illumina NovaSeq 6000 system (Illumina, USA) at Sequanta (China).

### 7. Alignment

For rat data, the FASTQ file was initially trimmed to remove Illumina adaptor sequences, and subsequently aligned with the Rattus norvegicus reference genome (NCBI GCF_015227675.2, Genome assembly mRatBN7.2) using HISAT2 (default parameters). This process generated a raw gene expression file.

For mouse data, the FASTQ file was initially trimmed to remove Illumina adaptor sequences, and subsequently aligned with the C57BL/6J Mouse reference genome (NCBI GCF_000001635.27,

Genome assembly GRCm39) using HISAT2 (default parameters). This process generated a raw gene expression file.

### 8. Reads Normalization

To minimize differences in sequence depth across various samples, normalization was performed as follows:

(1) Calculate the total reads for each sample.
(2) Adjust the total reads to 1 million reads for each sample.
(3) Back-calculate the reads per gene in each sample.

Normalized reads are referred to as Reads Per Million total reads (RPM).

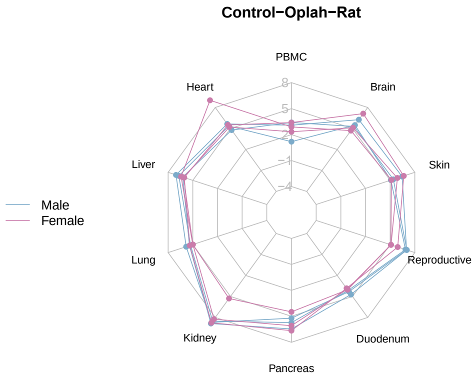


Fig. S3. Spider plot of *Oplah* count normalized tissue expression in healthy rat organs.

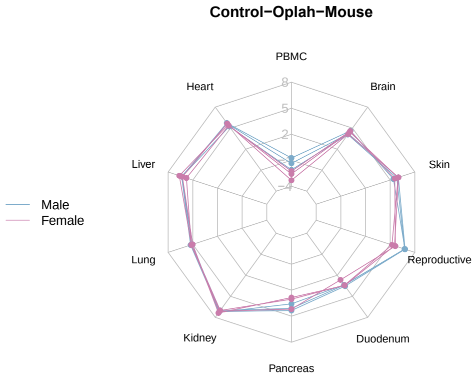


Fig. S4. Spider plot of *Oplah* count normalized tissue expression in healthy mouse organs.

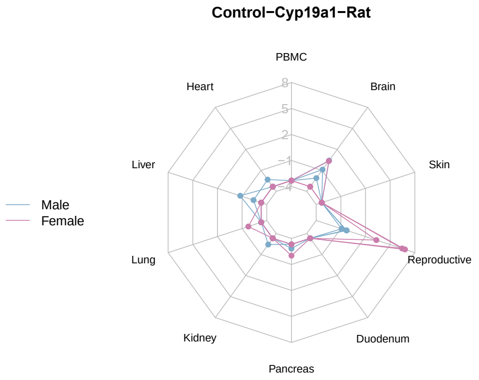


Fig. S5. Spider plot of *Cyp19a1* count normalized tissue expression in healthy rat organs.

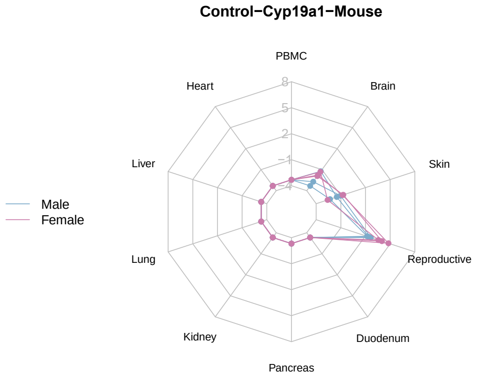


Fig. S6. Spider plot of *Cyp19a1* count normalized tissue expression in healthy mouse organs.

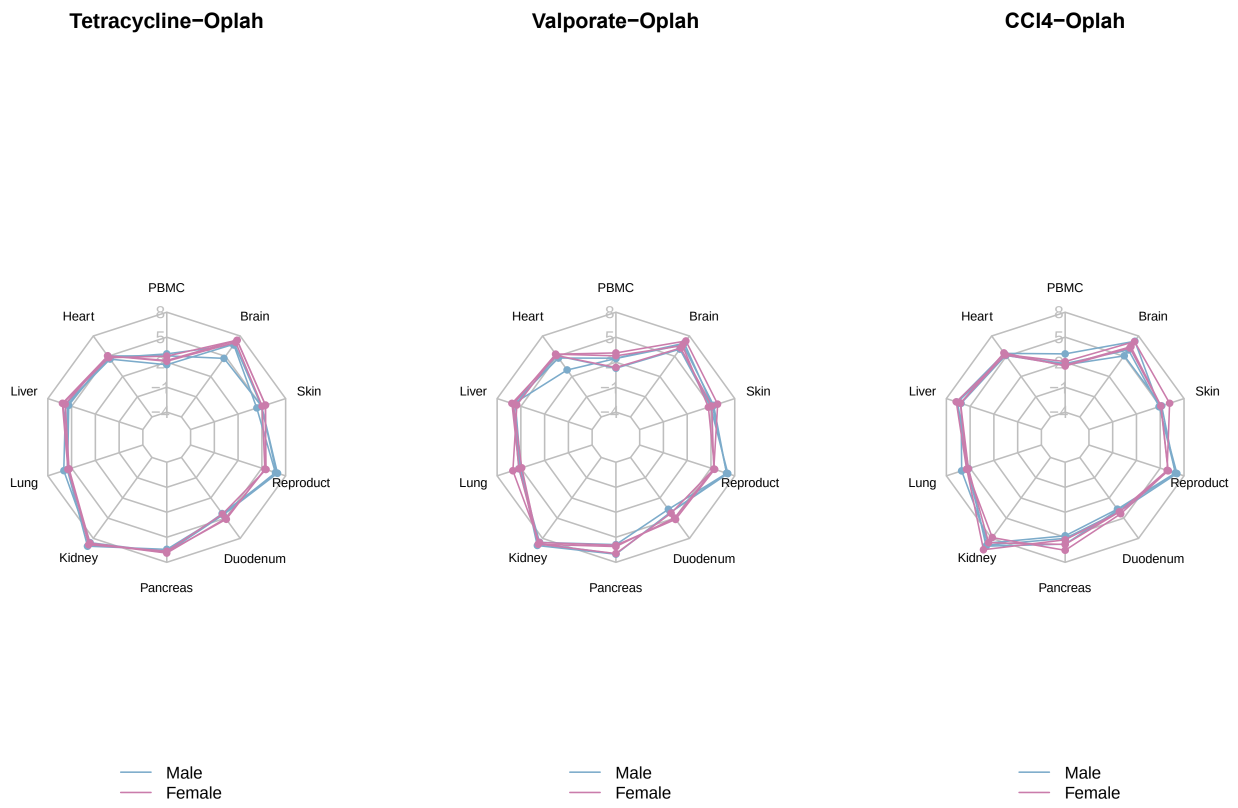


Fig. S7. Spider plot of *Oplah* reactome of rat organs post tetracycline, valproate, and CCl4 dosing.

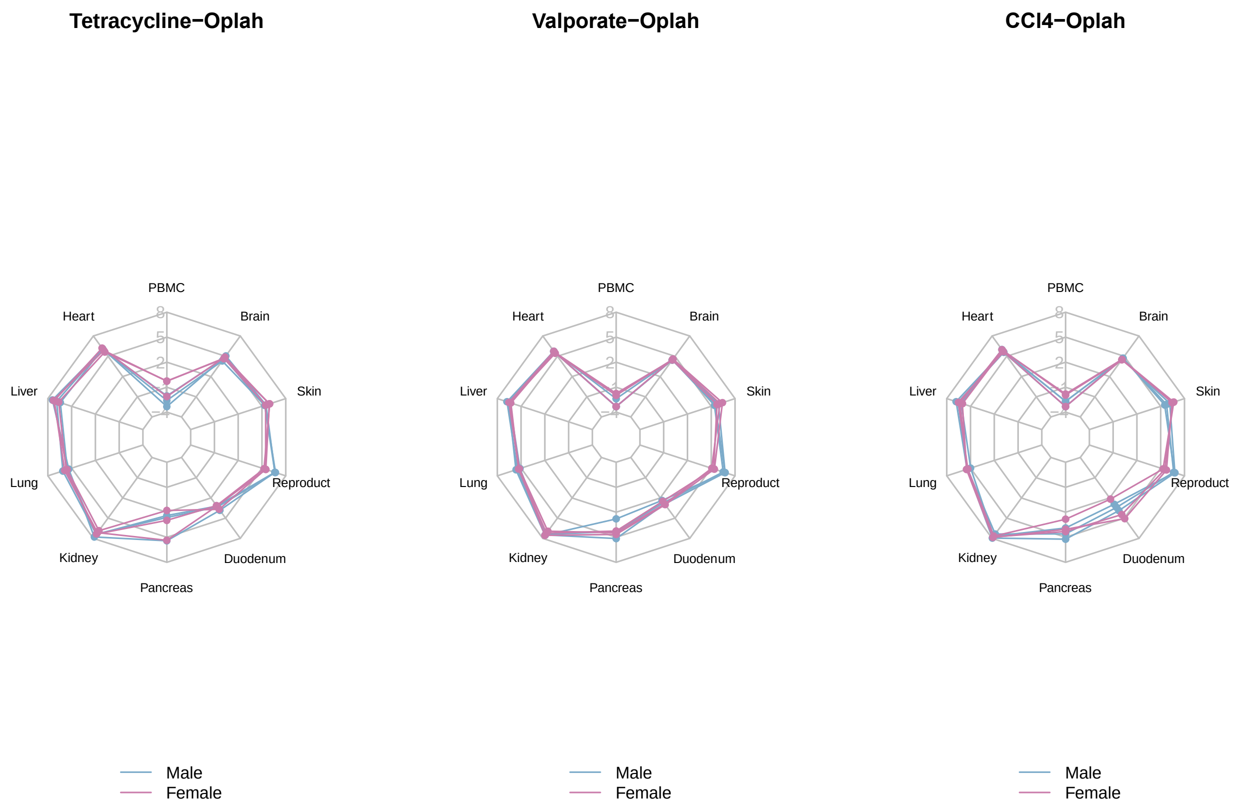


Fig. S8. Spider plot of *Oplah* count normalized tissue expression in mouse organs post tetracycline, valproate, and CCl4 dosing.

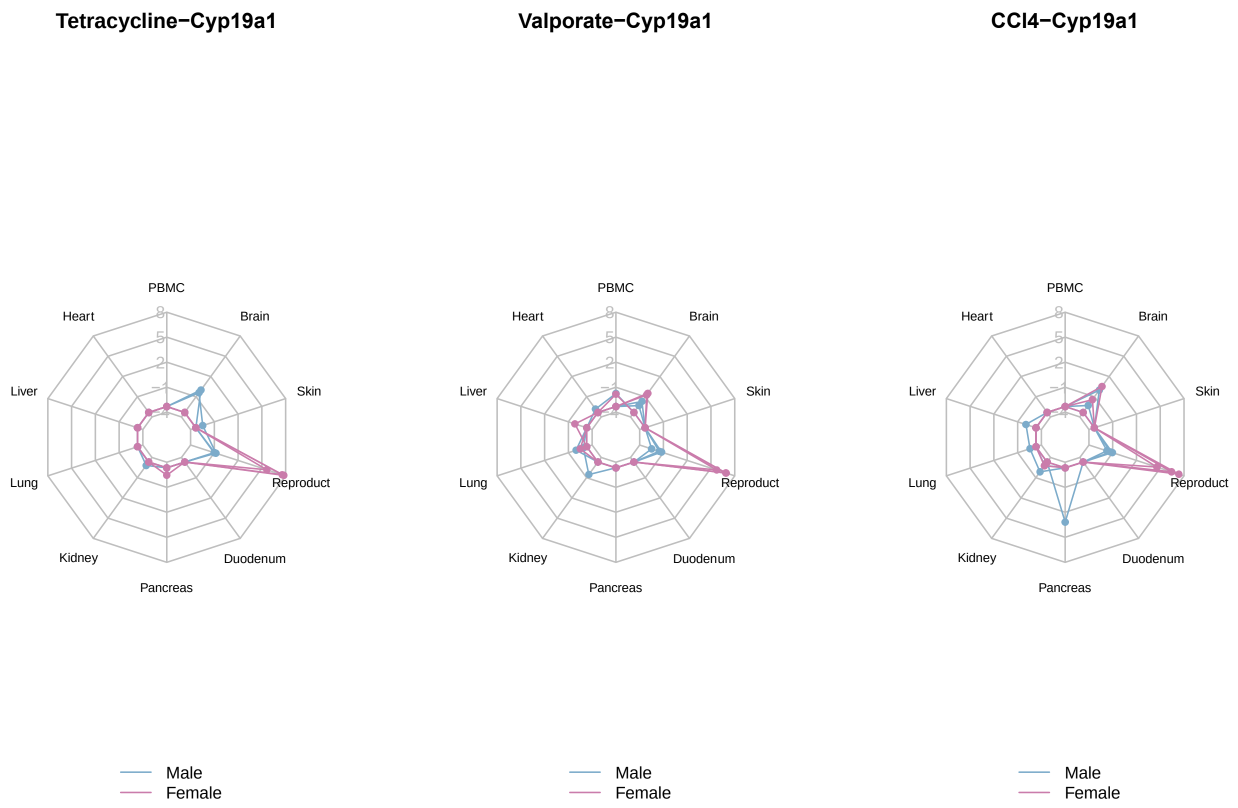


Fig. S9. Spider plot of *Cyp19a1* count normalized tissue expression in rat organs post tetracycline, valproate, and CCl4 dosing.

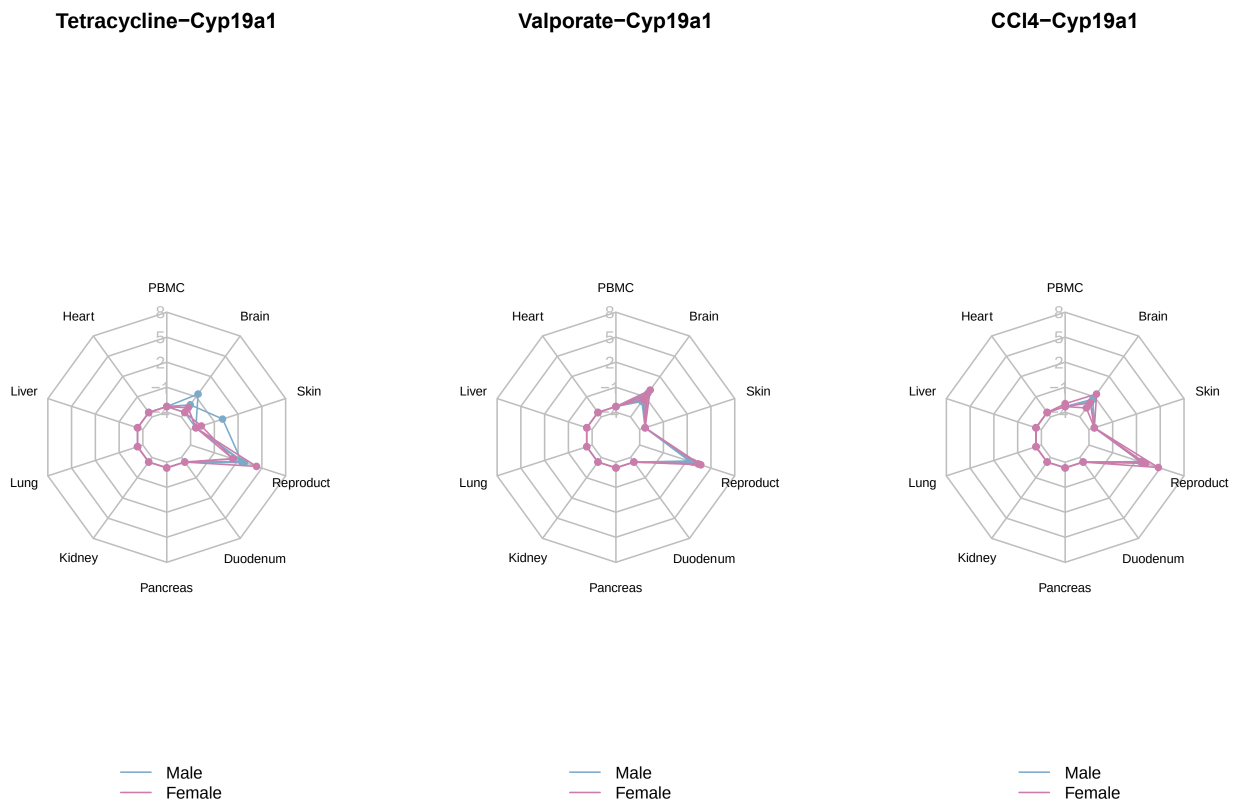


Fig. S10. Spider plot of *Cyp19a1* count normalized tissue expression in mouse organs post tetracycline, valproate, and CCl4 dosing.

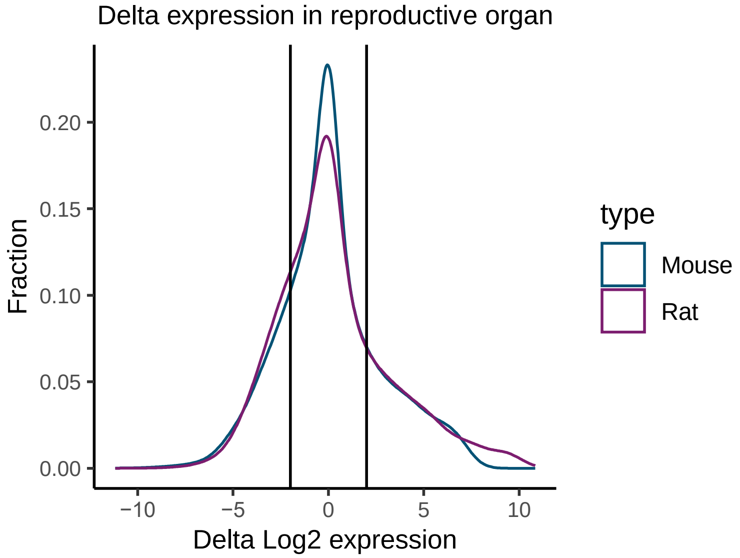


Fig. S11. Delta expression in reproductive organ for mouse and rat.

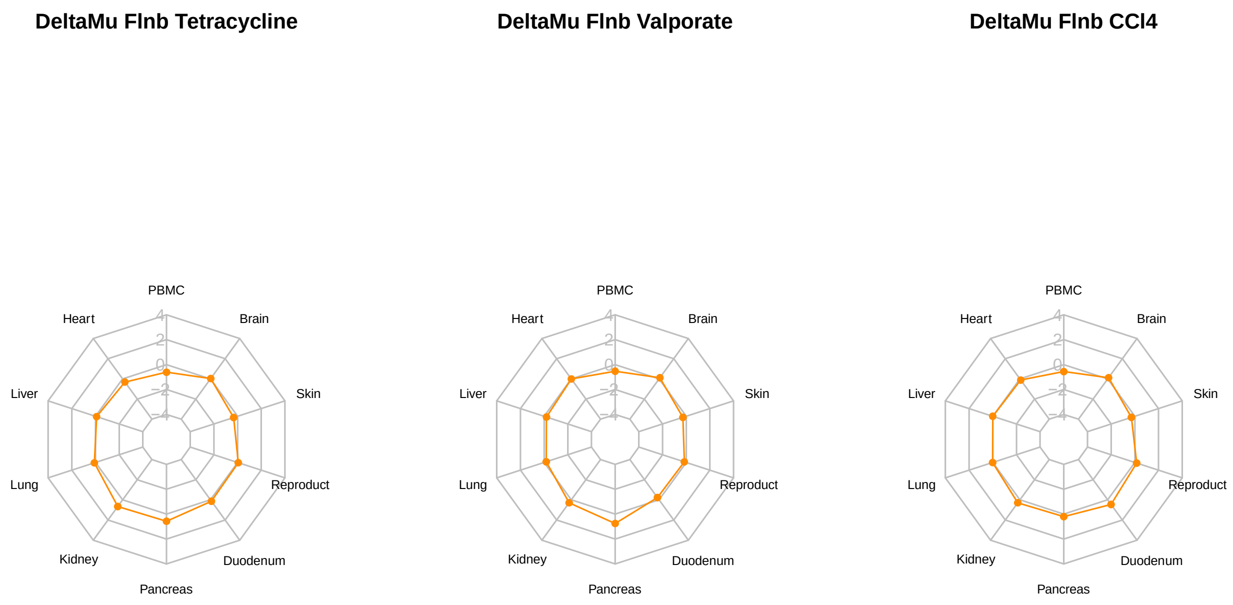


Fig. S13. Spider plot of *Flnb* reactome in rat organs post tetracycline, valproate, and CCl4 dosing.

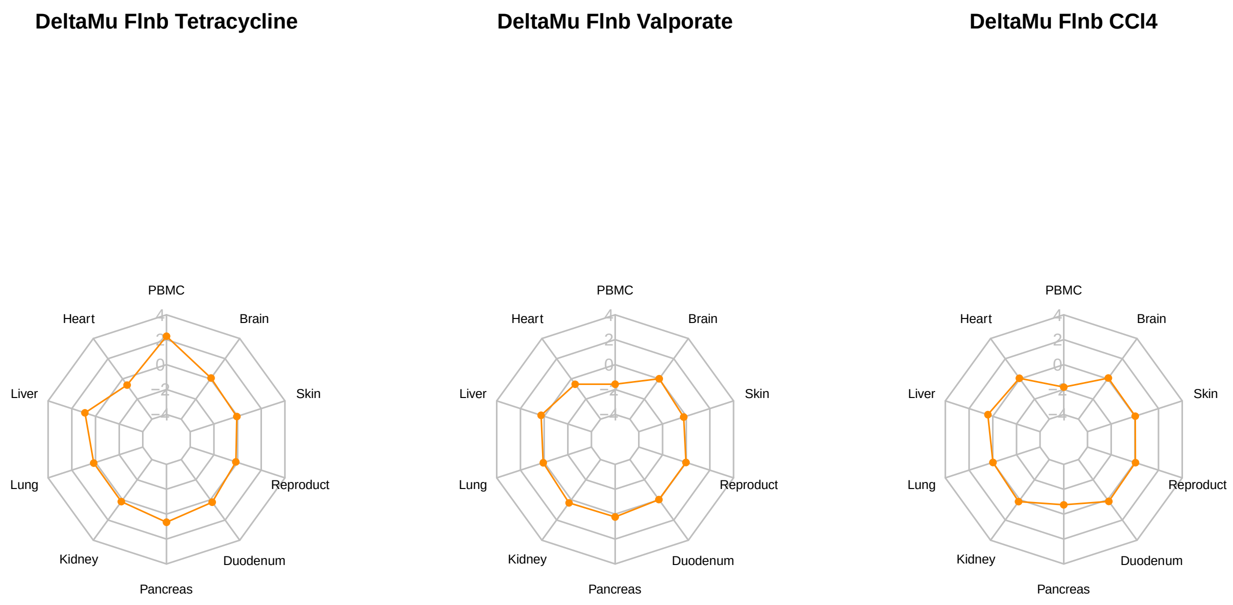


Fig. S14. Spider plot of *Flnb* reactome in mouse organs post tetracycline, valproate, and CCl4 dosing.

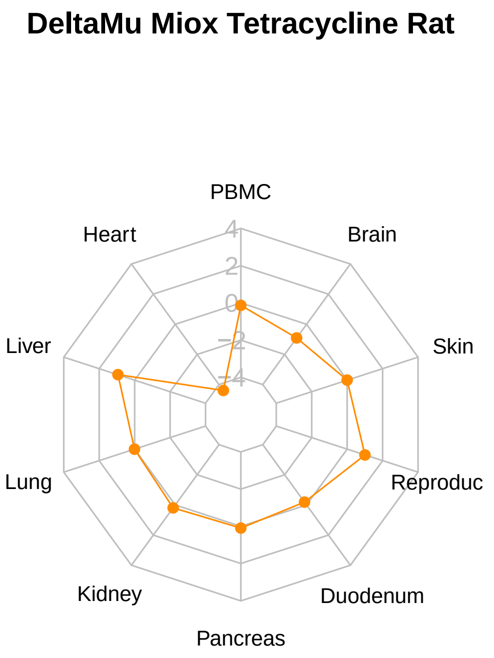


Fig. S15. Spider plot comparison of *Miox* reactome in mouse vs. rat organs post tetracycline dosing.

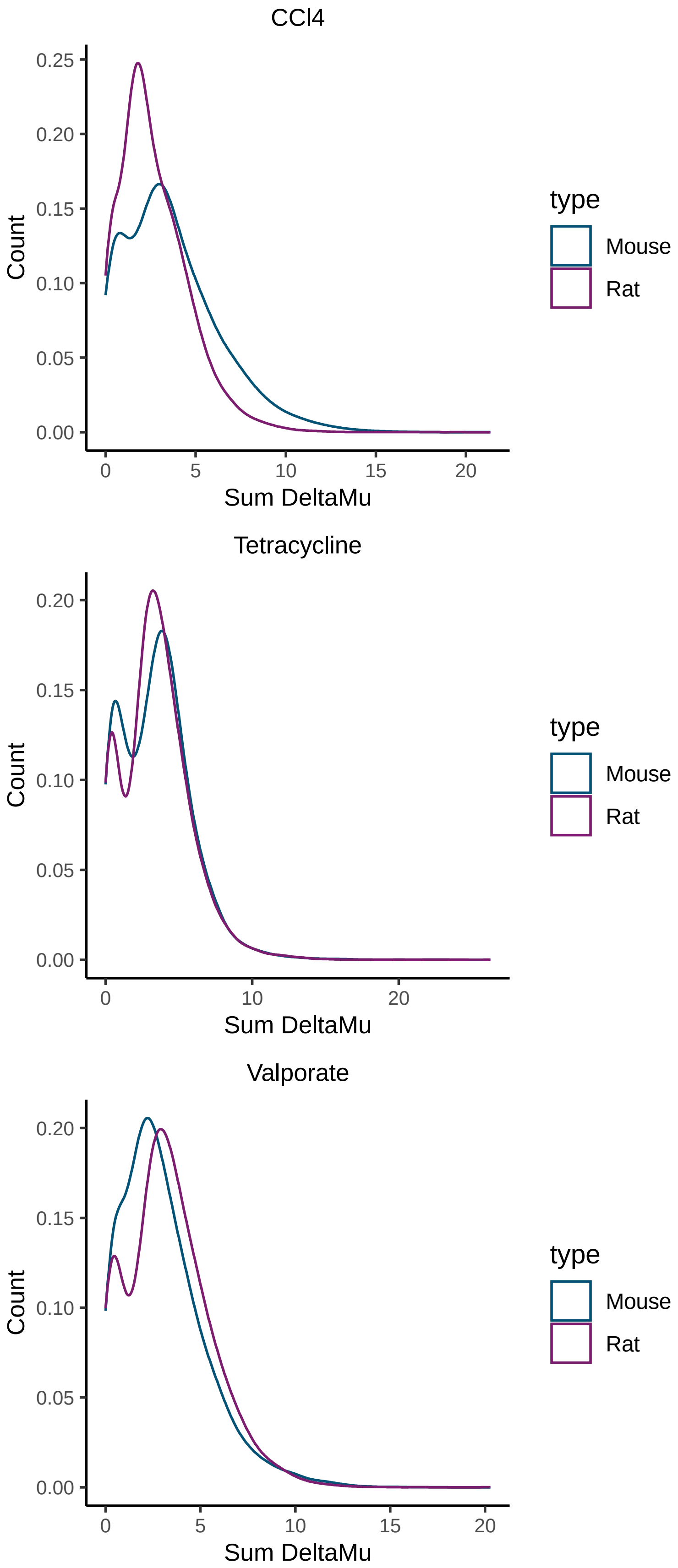


Fig. S16. Density plot of summed reactome value across organs in mouse vs. rat post tetracycline, valproate, and CCl4 dosing.

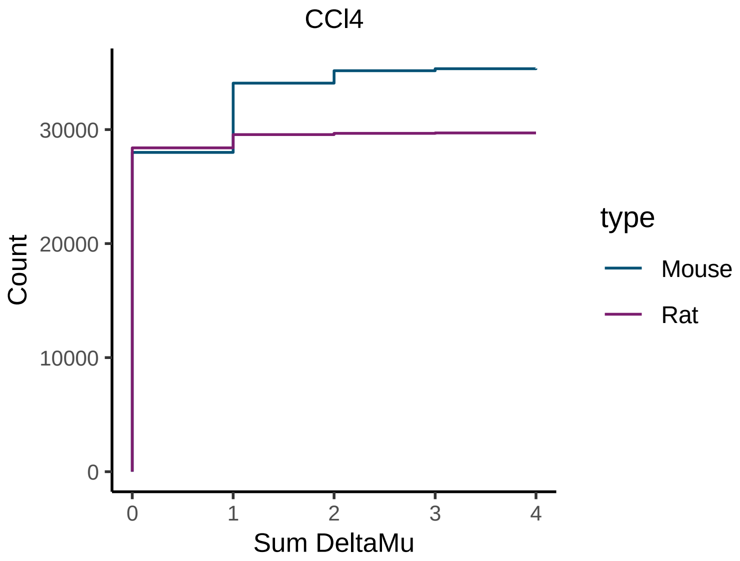


Fig. S17. Cumulative distribution function plot of summed hamming distance across organs in mouse vs. rat post CCl4 dosing.

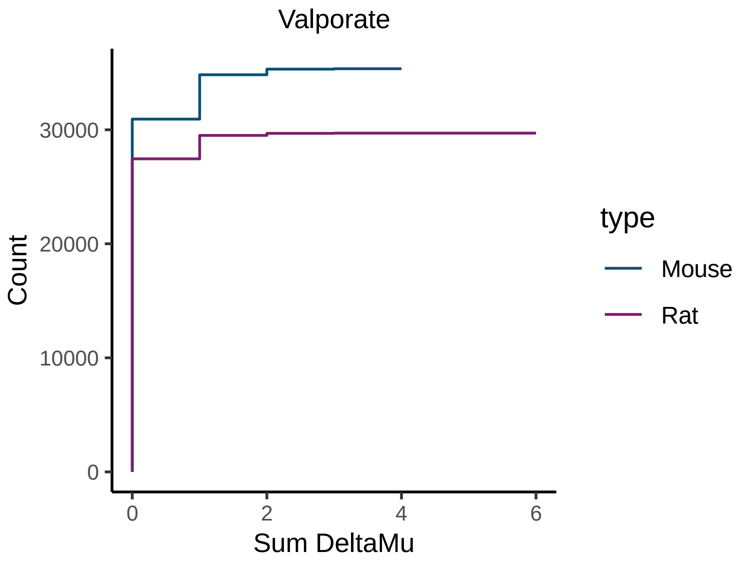


Fig. S18. Cumulative distribution function plot of summed hamming distance across organs in mouse vs. rat valproate dosing.

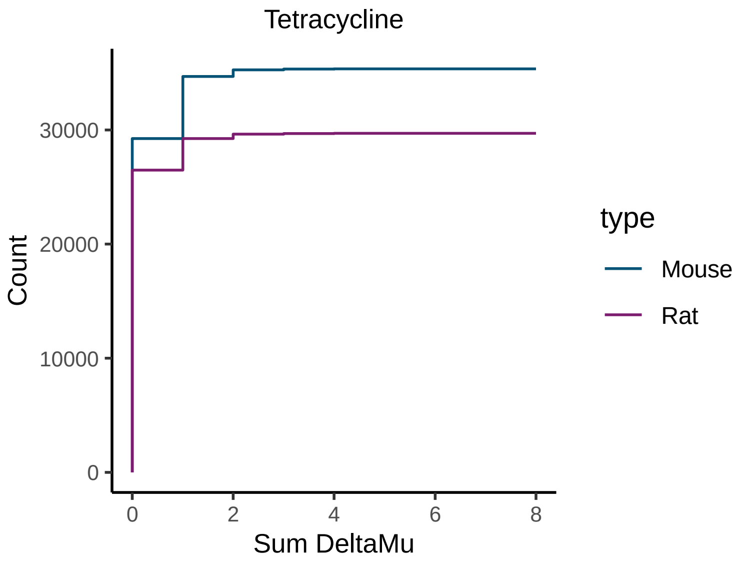


Fig. S19. Cumulative distribution function plot of summed hamming distance across organs in mouse vs. rat post tetracycline dosing.

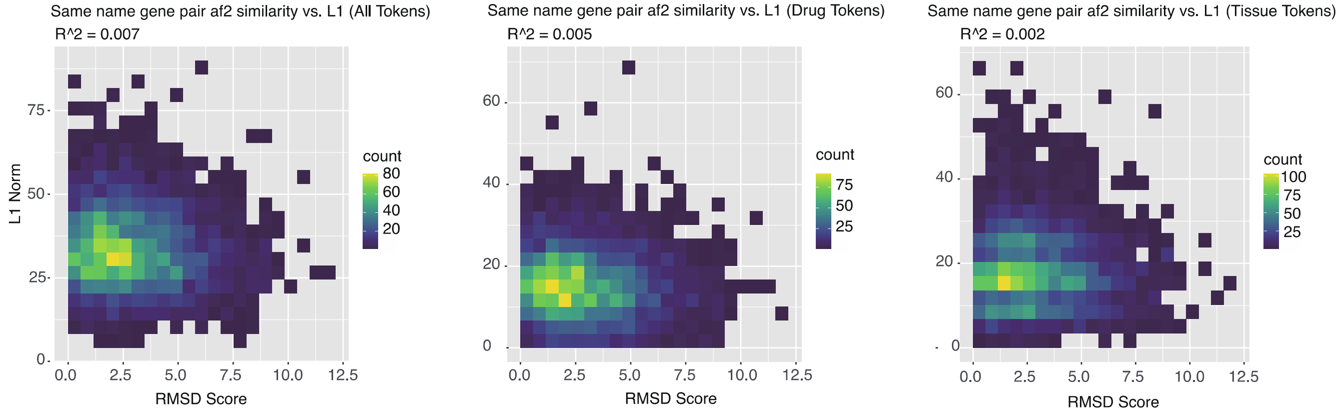


Fig. S20. correlation of Hamming distance based on all tokens (left), drug tokens (middle), and tissue tokens (right); y-axis: hamming distance calculated on L1 Norm, x-axis: Root Mean Square Error (RMSE) for average distance between the atoms of alphafold2 predicted protein structure for same name gene in mouse and rat. $R^2$ is calculated based on linear regression.

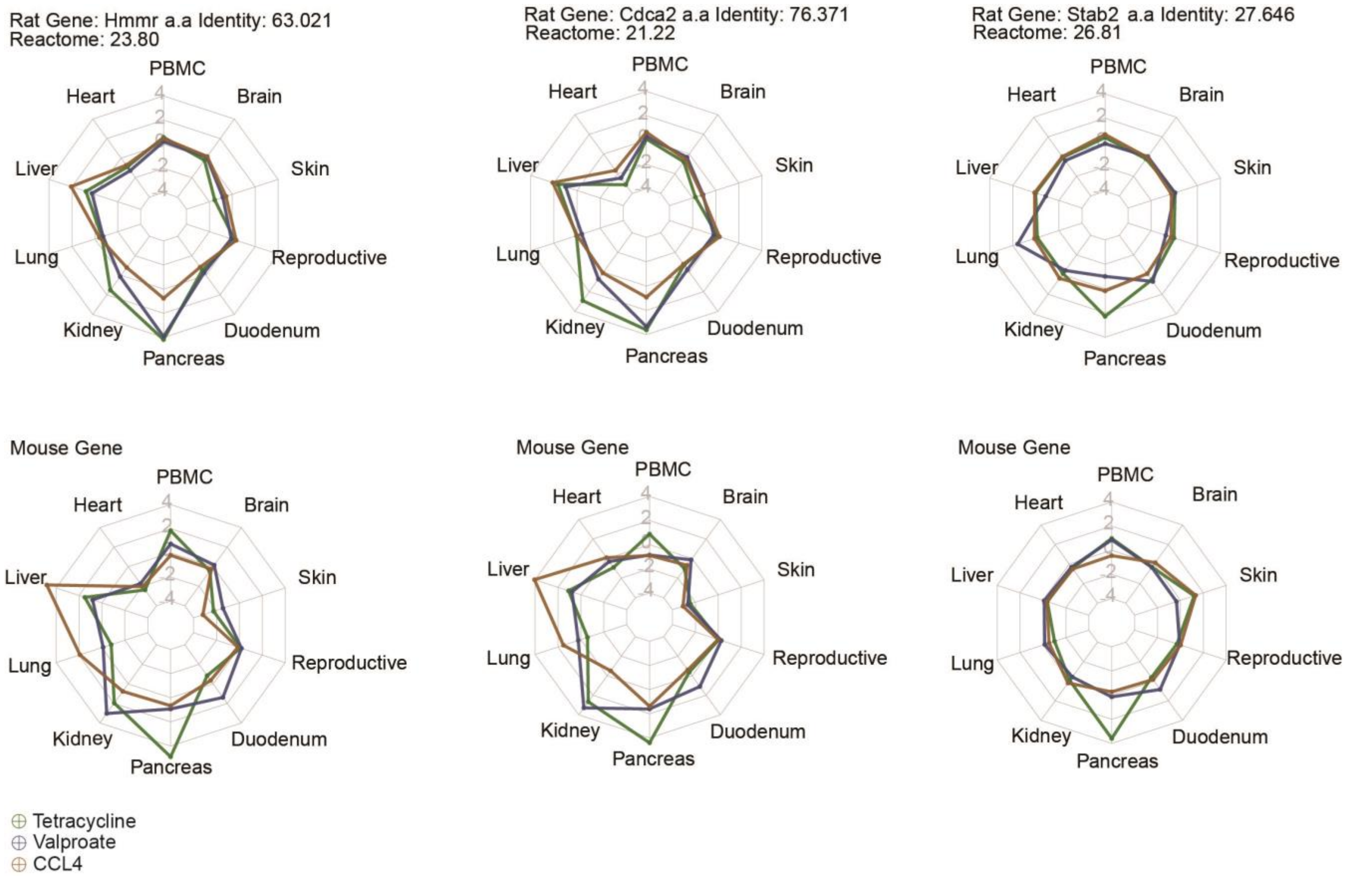


Fig. S21. More examples of Low Homology, Low Hamming Distance same-name gene pairs.

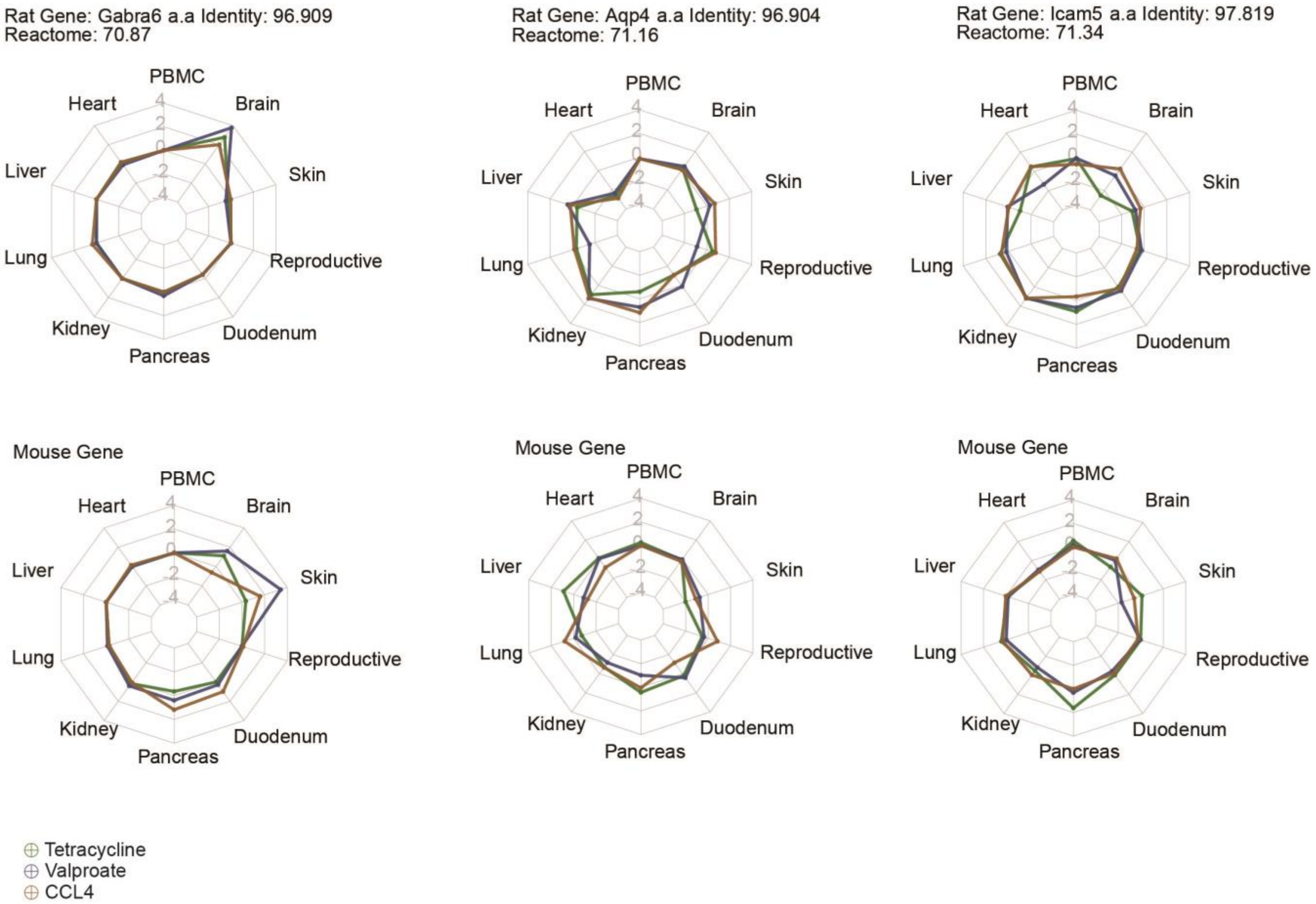


Fig. S22. More examples of High Homology, High Hamming Distance same-name gene pairs.

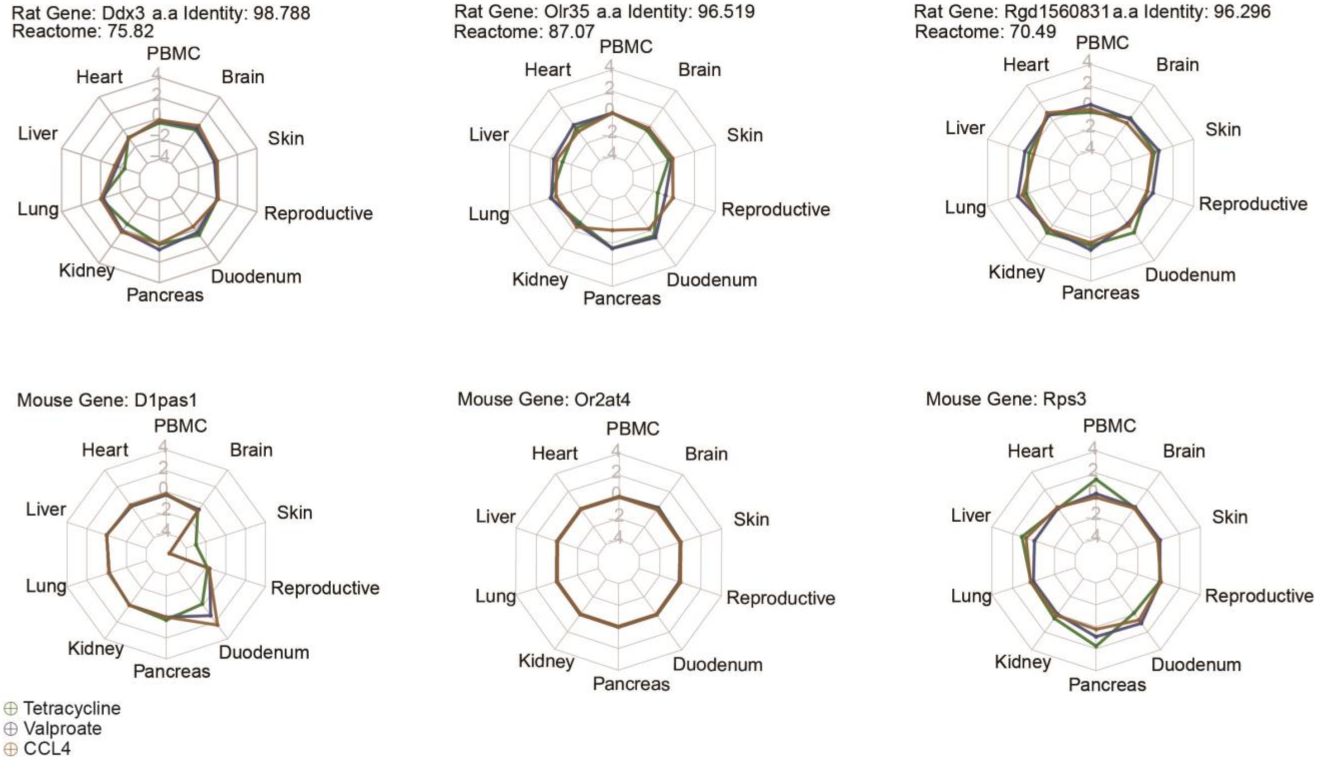


Fig. S23. Examples of High Homology, High Hamming Distance gene pairs (non-name matched).

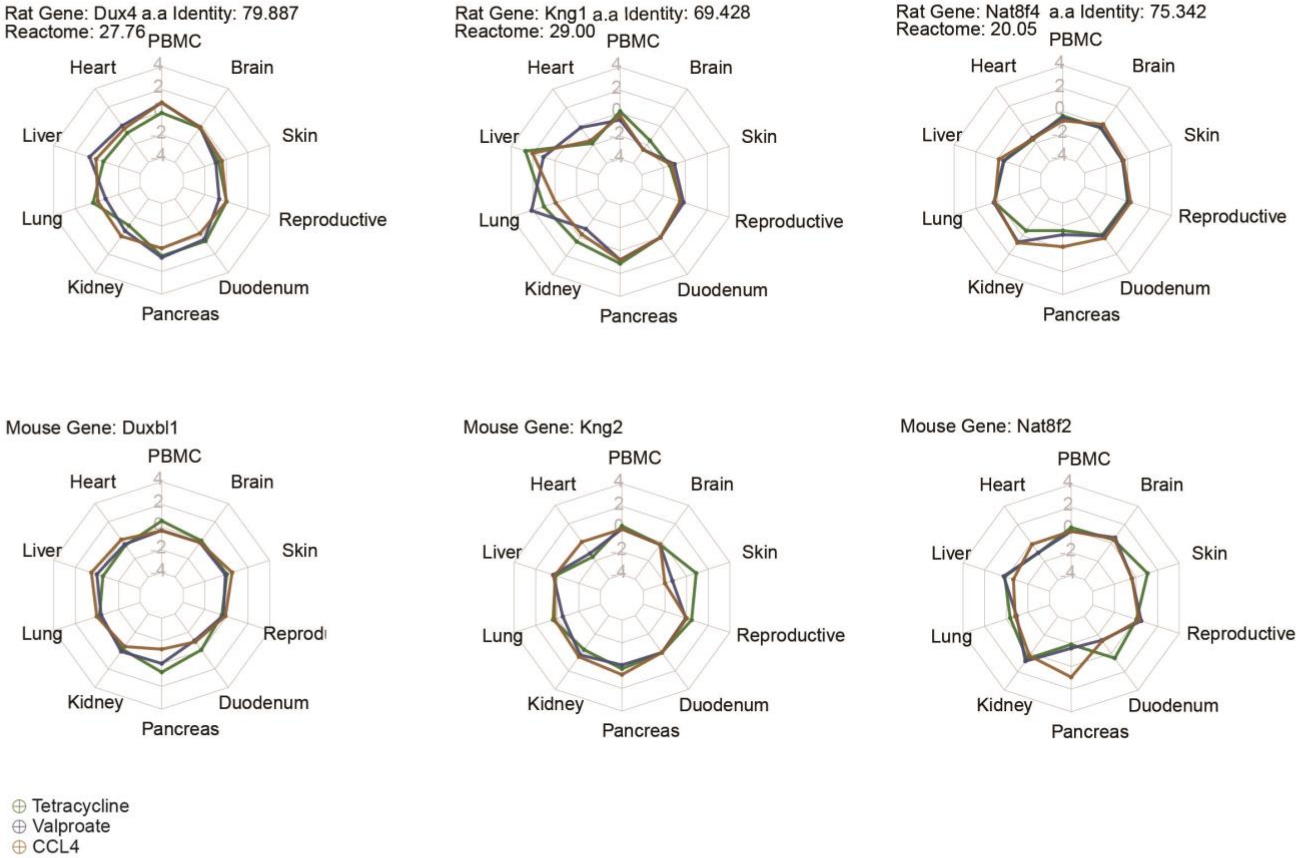


Fig. S24. Examples of Low Homology, Low Hamming Distance gene pairs (non-name matched).

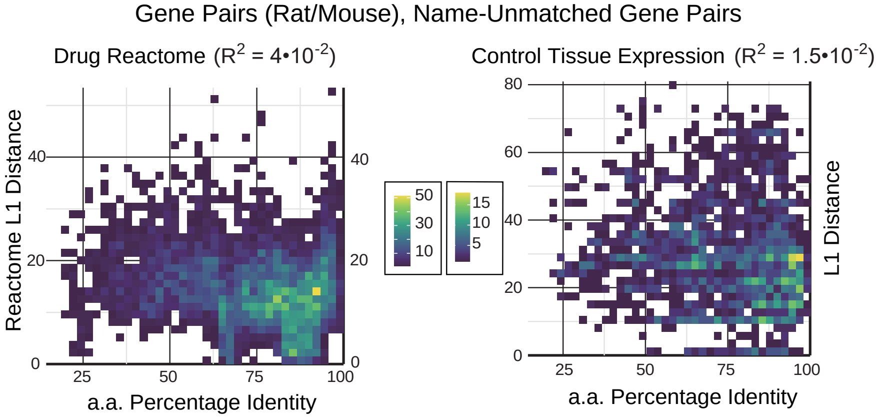


Fig. S25. Correlation of mouse and rat gene pairs based on best reactome (lowest hamming distance) matches. X- axis: amino acid sequence percentage identity based on blast; y-axis reactome hamming distance calculated on L1 Norm.

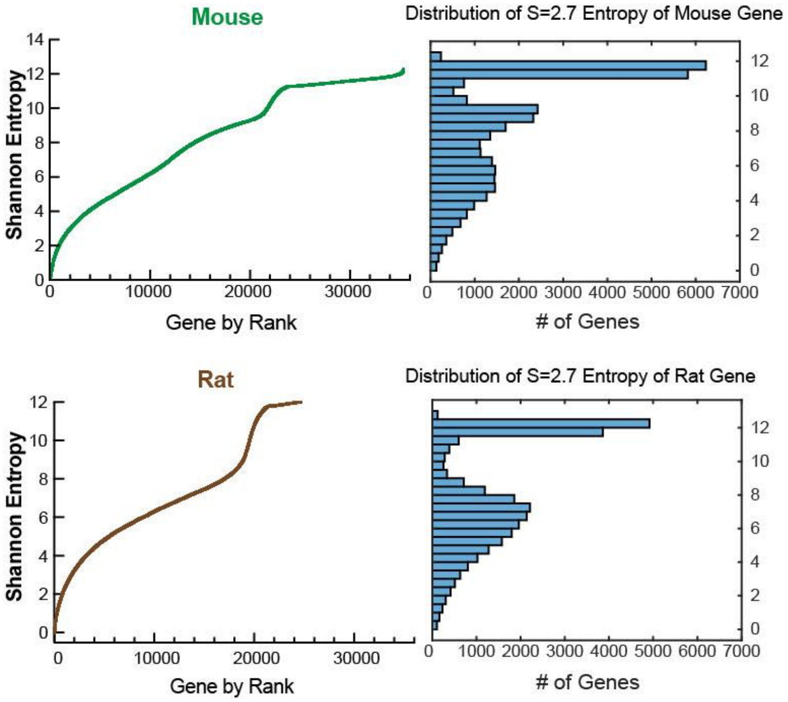


Fig. S26. Distribution of Shannon Entropies for mouse and rat genes, using both control tissue expression and drug reactomes, sorted in ascending order. Sharpness = 2.7.

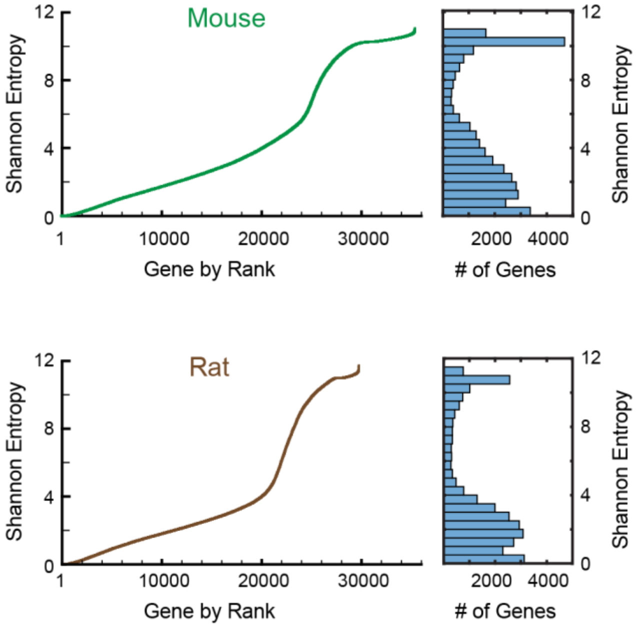


Fig. S27. Distribution of Shannon Entropies for mouse and rat genes, using both control tissue expression and drug reactomes, sorted in ascending order. Sharpness = 10.

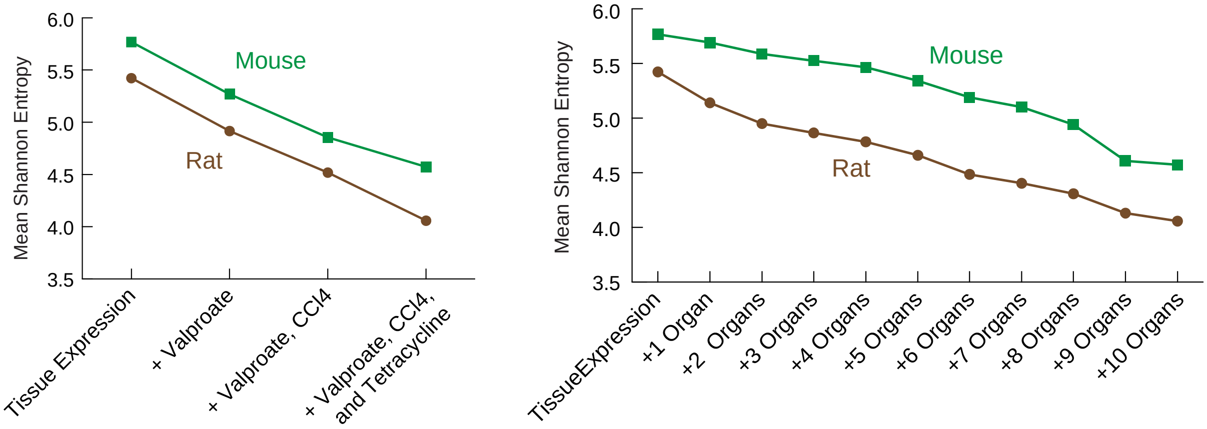


Fig. S28. Scaling properties of Shannon Entropy as additional drug reactome data or additional organ reactome data are included. Sharpness = 10.

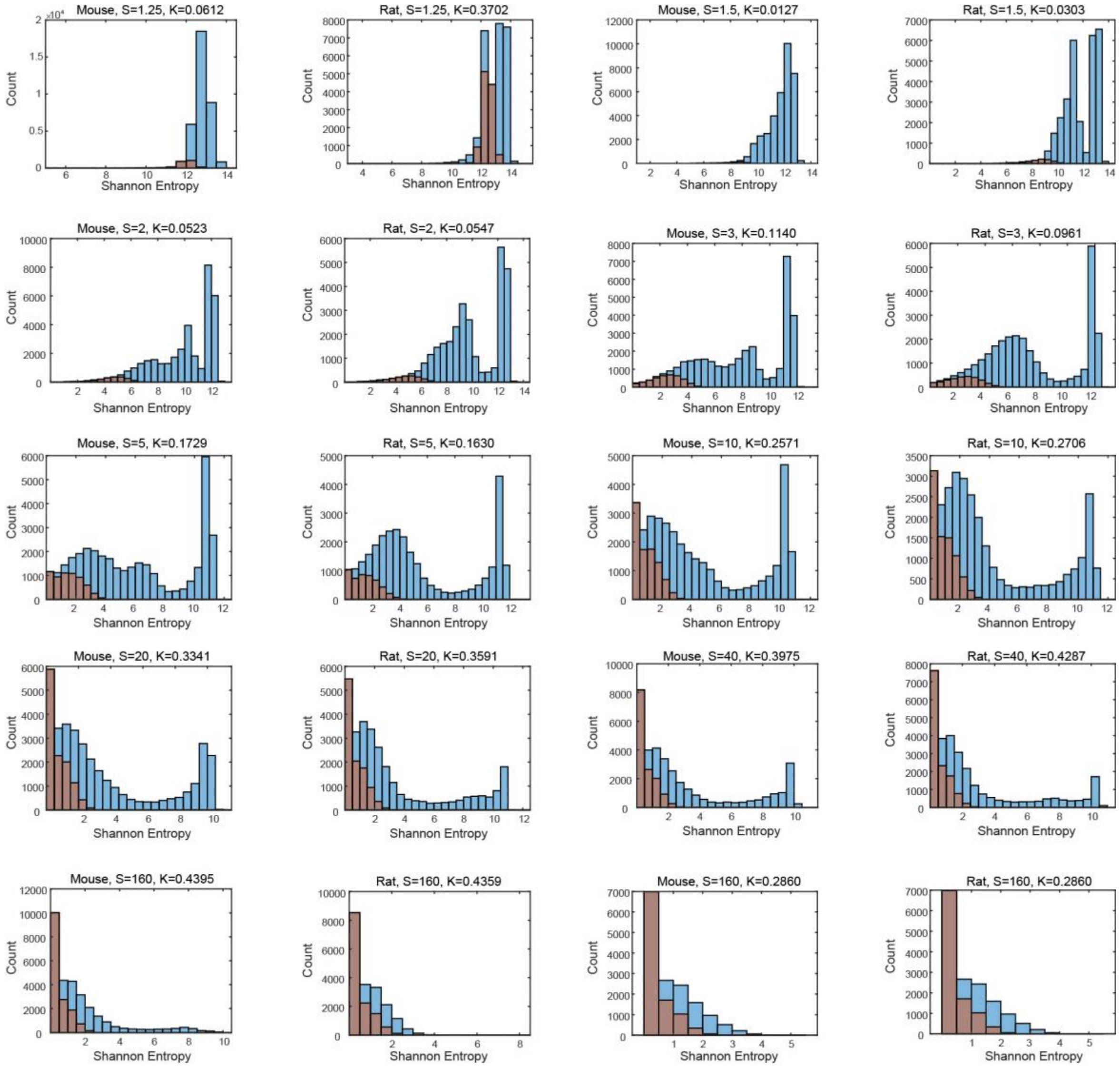


Fig. S29. Shannon Entropy distribution and value of K are observed for mouse=>rat and rat=>mouse gene mapping.
